# Insights into *Treponema pallidum* genomics from modern and ancient genomes using a novel mapping strategy

**DOI:** 10.1101/2023.02.08.526988

**Authors:** Marta Pla-Díaz, Gülfirde Akgül, Martyna Molak, Louis du Plessis, Hanna Panagiotopoulou, Karolina Doan, Wiesław Bogdanowicz, Paweł Dąbrowski, Maciej Oziębłowski, Barbara Kwiatkowska, Jacek Szczurowski, Joanna Grzelak, Natasha Arora, Kerttu Majander, Fernando González-Candelas, Verena J. Schuenemann

**Author notes:** Marta Pla-Díaz, Gülfirde Akgül, Martyna Molak, Louis du Plessis, Hanna Panagiotopoulou, Karolina Doan, Wiesław Bogdanowicz, Paweł Dąbrowski, Maciej Oziębłowski, Barbara Kwiatkowska, Jacek Szczurowski, Joanna Grzelak, Natasha Arora. These authors contributed equally to this manuscript (Marta Pla-Díaz, Gülfirde Akgül). Authors for correspondence: Verena J. Schuenemann < >, Kerttu Majander < >, Fernando González-Candelas < >.

## Abstract

**Background:** Treponemal diseases pose significant global health risks, presenting severe challenges to public health due to their serious health impacts if left untreated. Despite numerous genomic studies on *Treponema pallidum* and the known possible biases introduced by the choice of the reference genome used for mapping, few investigations have addressed how these biases affect phylogenetic and evolutionary analysis of these bacteria. In this study, we assessed the impact of selecting an appropriate genomic reference on phylogenetic and evolutionary analyses of *T. pallidum*.

**Results:** We designed a multiple-reference-based (MRB) mapping strategy using four different reference genomes and compared it to traditional single-reference mapping. To conduct this comparison, we created a genomic dataset comprising 77 modern and ancient genomes from the three subspecies of *T. pallidum*, including a newly sequenced 17th-century genome (35X coverage) of a syphilis-causing strain (designated as W86). Our findings show that recombination detection was consistent across different references, but the choice of reference significantly affected ancient genome reconstruction and phylogenetic inferences. The high-coverage W86 genome obtained here also provided a new calibration point for Bayesian molecular clock dating, improving the reconstruction of the evolutionary history of treponemal diseases. Additionally, we identified novel recombination events, positive selection targets, and refined dating estimates for key events in the species’ history.

**Conclusions:** This study highlights the importance of considering methodological implications and reference genome bias in High-Throughput Sequencing-based whole-genome analysis of *T. pallidum*, especially of ancient or low-coverage samples, contributing to a deeper understanding of this pathogen and its subspecies.

## BACKGROUND

Treponemal diseases such as syphilis, caused by *Treponema pallidum* subsp. *pallidum* (TPA), present persistent global health risks and can lead to severe health issues if left untreated (1,2). Historically, syphilis, mainly transmitted through sexual contact, has caused global epidemics since the end of the 15th-century until the era of effective antibiotic treatment. Currently, it is re-emerging worldwide in human populations. Two closely related treponematoses commonly transmitted through skin contact, yaws (caused by *T. pallidum* subsp. *pertenue*, TPE) and bejel (caused by *T. pallidum* subsp. *endemicum*, TEN), persist in developing countries (3,4). Yaws predominantly affects children (3,5–7) while there is limited epidemiological data available for bejel. However, bejel appears to be resurging in an unexpected clinical context (8,9).

Advances in high-throughput sequencing (HTS) technologies have enabled hundreds of new *T. pallidum* genomes to be published in recent years (10–23). Furthermore, several ancient genomes of this bacterium (24–27) have been reconstructed, using DNA extracted from archaeological remains of disease-causing organisms, a possibility previously inconceivable. This significant progress has provided detailed insights into the genomics of *T. pallidum* and facilitated vital epidemiological and evolutionary studies due to the significant incidence of treponemal diseases. Despite these developments, acquiring *T. pallidum* genomes remains a costly and labor-intensive procedure. Although an *in vitro* culture system exists (28,29), there is no standardized version applicable to all *T. pallidum* subspecies. Consequently, an enrichment process for the scarce DNA in clinical samples is still required.

Two primary methodological strategies are typically used to reconstruct individual pathogen genomes from the raw data from HTS: mapping to a reference genome and *de novo* assembly. However, achieving high-quality sequencing results necessary for *de novo* assembly can be challenging, even with modern samples. Consequently, mapping emerges as the predominant strategy for processing sequencing data of *T. pallidum* and obtaining the final modern genome sequences. While it has been possible to obtain ancient genomes from other pathogens through *de novo* assembly (30–35), the fragmented and degraded nature of ancient *T. pallidum* DNA presents significant challenges. Due to the low quantity of *T. pallidum* DNA and the presence of a wide range of other microbial DNA in the samples, no ancient *T. pallidum* genome has so far been obtained using this strategy. Nonetheless, well-characterized reference genomes and bioinformatic tools specifically designed for ancient data analysis, such as EAGER pipeline (36), are now enabling relatively efficient mapping for ancient treponemal genomes.

It should be noted that choosing the mapping strategy requires selecting the most closely related genome reference to the data being analyzed (37). Here, the best *T. pallidum* reference genomes among the known lineages were considered to be Nichols (CP004010.2) from TPA (Nichols lineage), SS14 (NC_021508.1) from TPA (SS14 lineage), CDC2 (CP002375.1) from TPE, and BosniaA (CP007548.1) from TEN, as these strains represent complete genomes obtained by *de novo* assembly (not partial or draft genomes), with minimal missing data (172 Ns out of 1.1Mb at most) (38). Moreover, the gaps between the contigs obtained from these samples have been closed by PCR and subsequent sequencing with Sanger technology. However, a recent study revealed point mutations in some of these genomic assemblies, resulting from successive passages in rabbits aimed at amplifying the DNA quantity for successful sequencing (29). The impact of these mutations requires further investigation, highlighting the need for careful consideration of the use of these reference genomes for mapping.

Despite the increase in genomic studies and the acknowledged potential for reference bias (37), few studies have reported detailed comparisons on the effects of reference selection (25,39,40). The investigations that have made such comparisons using different *T. pallidum* genomic references generally concluded that the differences were not significant and did not affect the conclusions, regardless of the chosen reference. However, these studies are constrained by the limited or unequal availability of genomic data across the subspecies and strains, particularly for TPE and TEN. Furthermore, no publications have considered the effect of reference selection on ancient genomes, which are by default more sparsely covered and possibly more derived, making them likely more vulnerable to these biases.

In this study, we explore the impact of the choice of genomic reference on the phylogenetic and evolutionary analyses of *T. pallidum* across its different subspecies, and in the context of an authentic ancient genome, reconstructed from a 17th-century strain (W86) at 35X coverage. Using a multiple-reference-based (MRB) mapping strategy with four reference genomes and traditional single-reference mapping, we examine 77 *T. pallidum* genomes, including the ancient W86 sample. Our findings reveal consistent recombination detection across diverse references and highlight the profound effect of reference choice on ancient genome reconstruction and phylogenetic interpretations. We also identify novel recombination events, positive selection targets, and refined dating estimates for key evolutionary events. Addressing these methodological implications and reference genome biases is crucial for advancing HTS-based whole-genome analysis of *T. pallidum*.

## RESULTS

### Pathogen screening for the new historical sample W86

A tooth sample yielding the new treponemal genome for this study was collected from individual W86, from the 17th century Ostrów Tumski cemetery in Wroclaw, Poland, documented to date from between 1621 and 1670 (41). The sample did not display paleopathological signs of infection and the genetic signal for *T. pallidum* was detected through a routine PCR-based screening for the presence of a range of selected pathogens. More information about the archaeological context of the newly obtained historical sample, as well as its anthropological and chemical characterization are detailed in Supplementary Notes 1-2.

The W86 sample was subsequently subjected to a screening procedure using direct shotgun sequencing (25,42,43). This process resulted in 581 unique reads that mapped against the SS14 strain, used as a *T. pallidum* genome reference, confirming the sample as positive for *T. pallidum* DNA. The reads showed 20% deaminated bases at the 5′ ends and 12% at the 3′ ends, with an average fragment length of 68 bp, signaling authenticity of ancient DNA (44,45) (for the damage profile, see Supplementary Figure 1). Following these screening and authentication steps, genome-wide enrichment for *T. pallidum* DNA (25,42,43) and HTS were conducted, resulting in 87.8 million raw reads.

### Single-reference-based (SRB) genome datasets and new ancient genome reconstruction

We generated four single-reference-based (SRB) genome datasets by mapping each of the 77 genomes selected for this study against each of the four genomic references chosen as representatives of *T. pallidum* subspecies and/or clades (CDC2, BosniaA, Nichols, and SS14) (see Supplementary Table 1). We will denote these SRB mapping datasets as CDC2-SRB (Supplementary File 1), BosniaA-SRB (Supplementary file 2), SS14-SRB (Supplementary file 3), and Nichols-SRB (Supplementary file 4) datasets, respectively. We obtained the following lengths and number of SNPs for each one of these mapping-genome datasets: CDC2-SRB (1,139,744 bp, 3529 SNPs), BosniaA-SRB (1,137,653 bp, 3403 SNPs), SS14-SRB (1,139,569 bp, 3429 SNPs), and Nichols-SRB (1,139,633 bp, 3593 SNPs).

After generating the four SRB genome datasets, a maximum likelihood (ML) tree was constructed for each one. We then analyzed the newly acquired ancient genome, W86, to determine its position in each of the phylogenetic trees derived from the four SRB genome datasets. This analysis aimed to ascertain its classification within the three subspecies and/or clades of *T. pallidum*. The SS14 strain was identified as the most closely related reference genome to the ancient genome W86, leading to the classification of W86 as an SS14-like strain of *T. pallidum pallidum* (TPA) (see Supplementary figure 2).

The W86 sample produced 524,587 unique treponemal reads successfully mapped to the SS14 reference. This resulted in a comprehensive coverage of 98.21% of the bases by a minimum of 3 reads and a median depth coverage of 35X (Supplementary Table 1). The variant calling process revealed the presence of 163 SNPs with respect to the SS14 reference genome. In order to analyze the sensitivity of this new ancient genome to macrolide antibiotics, two known *T. pallidum* 23S ribosomal RNA gene mutations (A2058G and A2059G) were examined. However, neither of these mutations were present in the W86 genome, indicating that this strain is sensitive to macrolide antibiotics.

**Figure 1.**
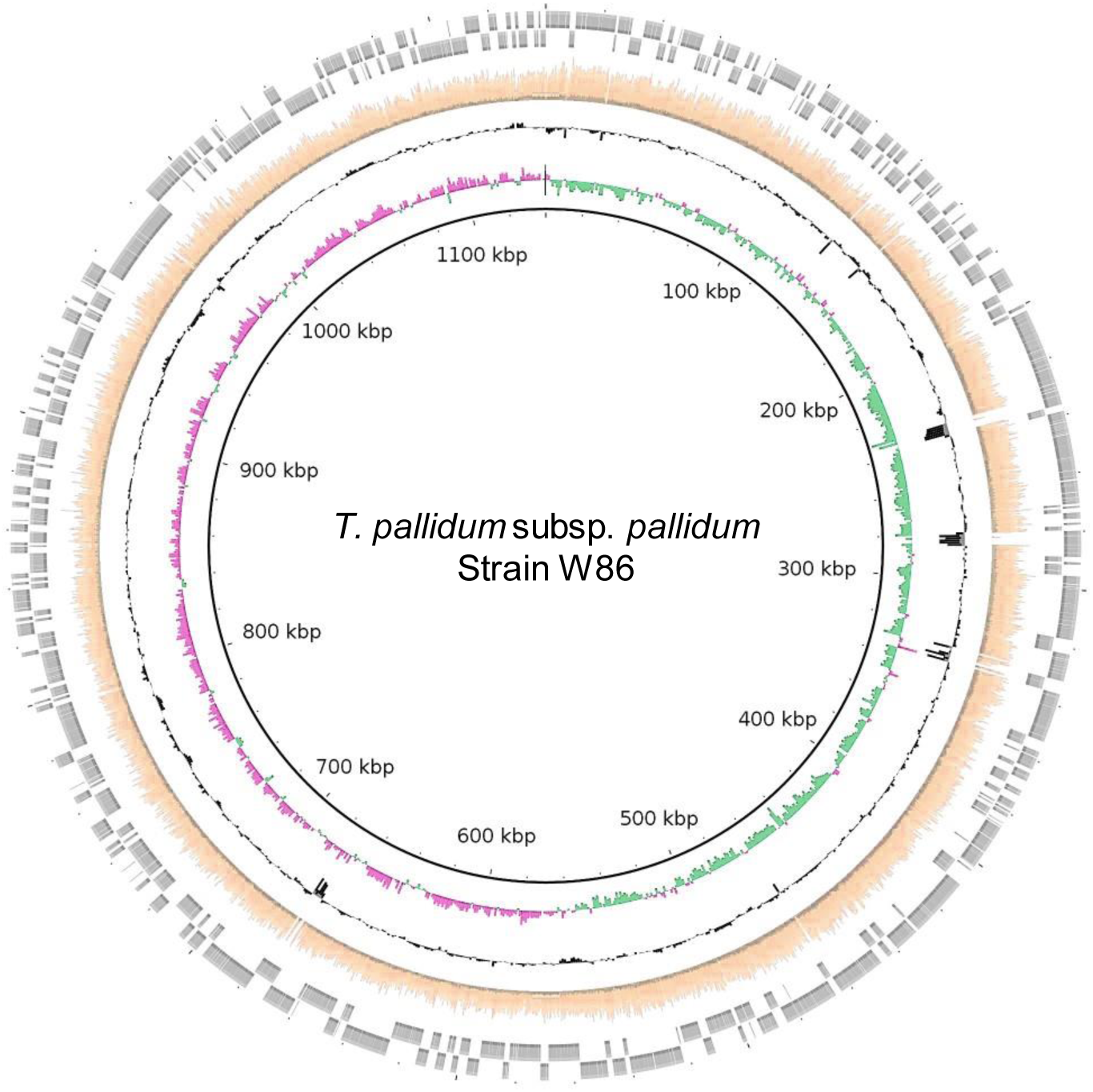
Circular plot of the W86 genome. Circles indicate, from inside outwards: genomic position, GC skew (pink and green); GC content (black) and coverage (orange). The outer rim (gray) shows the direction of protein-coding regions according to the annotation of the SS14 reference genome (CP000805.1): forward, outermost circle.

### Multiple-reference-based (MRB) alignment

A multiple-reference-based (MRB) genome alignment was created for the 61 *T. pallidum* strains from previous studies with available raw data, using their nearest reference genomes (Supplementary File 5). Furthermore, for the 15 strains lacking raw data, their genome assembly sequences were added to the previously obtained MRB alignment and realigned. This MRB genome alignment of 77 different genomes (including the newly acquired ancient genome, W86, whose sequence was selected from SS14-SRB) covers a total of 1,141,136 nucleotides (Supplementary File 3). In this alignment, 4,822 SNPs were identified. Detailed information on each of these genomes is available in Supplementary Table 1.

To detect orthologous genes in the four reference genomes employed and to identify the location of genes in the MRB genome alignment, we first conducted an orthology analysis with the four reference genomes (Supplementary Table 2). This analysis revealed a total of 1137 distinct orthology groups, 21 of which were found to have more than one gene per subspecies/sublineage, resulting in a total of 1158 unique genes. Additionally, some *tpr* genes (*tp0316, tp0317, tp0621, tp0620*) known to undergo gene conversion and to occupy different genomic locations among the four reference genomes were extracted separately, yielding a total of 1161 genes. The genomic coordinates for the individual 1161 genes in the MRB genome alignment and the SNPs detected in each of them are detailed in Supplementary Tables 3 and 4, respectively. We also generated a ML tree for the MRB alignment as we did for each one of the SRB genome datasets.

### MRB alignment comparison with SRB genome datasets

We assessed topological and evolutionary differences among the four SRB trees and the MRB tree by computing Robinson-Foulds (RF) distances with RAxML (Table 1). The average RF distances following pairwise tree comparisons were relatively similar to each other.

**Table 1.** Robinson-Foulds distance (RF) obtained by RAxML of the trees obtained from the distinct mapping datasets and the whole-genome phylogeny.

| Tree1 | Tree2 | Number of Branch changes | RF value | Average RF value |
| --- | --- | --- | --- | --- |
| BosniaA | CDC2 | 72 | 0.487 | 0.493 |
| BosniaA | Nichols | 64 | 0.432 |  |
| BosniaA | SS14 | 74 | 0.500 |  |
| BosniaA | MRB | 82 | 0.554 |  |
| CDC2 | Nichols | 70 | 0.473 | 0.483 |
| CDC2 | SS14 | 72 | 0.487 |  |
| CDC2 | MRB | 72 | 0.487 |  |
| Nichols | SS14 | 60 | 0.405 | 0.449 |
| Nichols | MRB | 72 | 0.487 |  |
| SS14 | MRB | 54 | 0.365 | 0.439 |
| MRB | - | - | - | 0.473 |

Furthermore, the four SRB trees obtained were visually compared with the MRB phylogeny to identify any topological differences (see Supplementary Figures 2-5).

Based on our findings, and building on previous phylogenetic classifications and nomenclature of the SS14 lineage (42,46), we defined the SS14-Ω sublineage as the clade that includes most of the SS14 genomes from clinical and modern samples, which was previously defined as a mostly epidemic, macrolide-resistant cluster that emerged after, and possibly prompted by, the discovery and widespread use of antibiotics (42,46). The SS14 lineage includes all ancient TPA genomes (including the new ancient genome W86) and the Mexico A strain, while the SS14-Ω sublineage encompasses all modern SS14 genomes from clinical samples.

The most relevant topological variations in the SRB trees, in comparison with the MRB tree, were found in the SRB trees generated from datasets where BosniaA, CDC2, and Nichols were used as reference genomes. These trees exhibited topological changes across all clades (TEN, TPE, Nichols, and SS14), with the most notable ones occurring in the Nichols and SS14 clades (Supplementary Figures 2-5).

The most remarkable differences affected the SS14 strains, especially the four ancient TPA genomes (including W86) and the Mexico A strain. Specifically, in the tree derived from the BosniaA-SRB genome dataset (Supplementary Figure 3B), the ancient genomes 94A and 94B were at the base of all TPA strains, and Mexico A was located within the SS14-Ω clade. This is in contrast to the MRB tree (Supplementary Figure 3A), in which both ancient genomes were basal to all SS14 strains. Similar topological incongruences were observed in the tree obtained using CDC2-SRB with regards to the 94A and 94B ancient genomes (Supplementary Figure 2B), although in this dataset, Mexico A remained basal to all SS14 strains.

However, in the tree obtained using the Nichols-SRB genome dataset (Supplementary Figure 4B), the most notable topological change occurred with the Mexico A strain. Similarly to the tree obtained from the CDC2-SRB dataset (Supplementary Figure 2B), Mexico A was located within the SS14-Ω clade and not basal to the entire SS14 clade. Unlike the ML trees obtained from the previously detailed SRB genome datasets (Supplementary Figures 2-4), the ancient strains 94A and 94B occupied the same position in the tree as in that obtained using the MRB genome alignment. Nevertheless, the ancient strain SJ219 appeared in the ML tree obtained with the Nichols-SRB dataset (Supplementary Figure 4B) with a much longer branch, implying many more inferred substitutions in this strain.

Moreover, in all four ML trees obtained with the SRB genome datasets (Supplementary Figures 2-5) remarkable changes were also observed in the Nichols clade when they were compared to the ML tree obtained with the MRB genome alignment. In the trees obtained using the BosniaA-, Nichols-, and SS14-SRB genome datasets (Supplementary Figures 3B-5B), the Seattle 81-4 strain, which was basal to all the strains of the Nichols clade in the MRB tree, grouped among other strains of the Nichols clade. Additionally, a remarkable change was observed regarding strain CW82. Instead of grouping with other Nichols strains as in the MRB tree, it occupied a basal position to the Nichols strains in all four of the ML trees obtained using the SRB genome datasets (Supplementary Figures 2B-5B).

### Analysis of recombination

To obtain a reliable phylogenetic reconstruction, it is necessary to remove genomic regions that are not strictly subject to vertical inheritance, e.g. recombinant regions or loci with intra- or intergenic conversion. Previous studies examining different sets of genomes have identified and analyzed such loci (25,38,40,42,47–49). Here, we proceeded to carry out a comprehensive investigation of recombination and its impact, using the MRB genome alignment generated. Our detailed recombination detection pipeline, the phylogenetic incongruence method (PIM), yielded 28 recombinant regions. These derived from 20 different genes, and encompassed a total of 1,114 SNPs (21.24% of the total SNPs) among the *T. pallidum* strains analyzed here (Table 2). The average length of the recombinant regions was 441 bp, with a minimum length of 4 bp and a maximum of 2,097 bp. Details on the intermediate results of the likelihood mapping and topology tests from the PIM procedure applied to these genomes are provided in the Supplementary Material (Supplementary Note 4 and Supplementary Tables 4-6).

**Table 2.** Recombination events detected in *T. pallidum*. The gene ID names correspond to the general gene nomenclature for *T. pallidum.* For each recombination event, coordinates for the start and end position in the multiple genome alignment are provided (Supplementary File 5). The strains involved are detailed, with an arrow separating the donor strains from the recipient strains. Events with an asterisk may represent more than one recombinant transfer depending on the placement of the involved strains in the reference tree.

| Gene ID | Event | Start | End | Minimum size (bp) | SNPs | Strains Involved |
| --- | --- | --- | --- | --- | --- | --- |
| *tp0131* | 1 | 152624 | 153390 | 766 | 333 | Gauthier, LMNP-1, CDC_2575 → Nichols, Chicago, BAL3,BAL73, NIC2 |
| *tp0136* | 1 | 158235 | 158247 | 12 | 4 | TPE → NL16 |
|  | 2* | 158281 | 158292 | 11 | 4 | Seattle 81-4 → CW82, CW83 |
|  | 3 | 158292 | 158310 | 18 | 4 | TPE clade, excluding HATO, OKA_2116 and LMNP-1 → Seattle 81-4 |
|  | 4 | 158414 | 158418 | 4 | 4 | TPE → Nichols-lineage clade excluding CW82 |
|  | 5 | 158483 | 158507 | 24 | 8 | TEN → Nichols-lineage clade |
|  | 6* | 159058 | 159119 | 61 | 4 | TEN/TPE → PD28 and Nichols-lineage clade excluding CW82 |
|  | 7 | 159452 | 159466 | 14 | 6 | TEN/TPE → Nichols-lineage clade |
| *tp0164* | 1* | 187135 | 187320 | 185 | 5 | TEN/TPE → Seattle 81-4, CW86, CW59, NE20, 94A, 94B |
| *tp0179* | 1* | 198183 | 198571 | 388 | 9 | TEN/TPE → Nichols-lineage clade, W86, PD28 |
| *t0012* | 1 | 233067 | 233223 | 71 | 33 | TEN/TPE excluding CDC_2575, SamoaD, K363, K403 → Mexico A |
| *t0015* | 1 | 281513 | 281670 | 74 | 48 | TEN/TPE excluding CDC_2575, SamoaD, K363, K403 → Mexico A |
| *tp0326* | 1 | 346656 | 348753 | 2097 | 59 | TEN → SS14-lineage clade, excluding Mexico A and syphilis ancient genomes |
| *tp0346* | 1* | 373266 | 373282 | 16 | 3 | TEN/TPE → 94A, 94B |
| *tp0462* | 1 | 493382 | 494407 | 1025 | 55 | TEN/TPE → CW86, Seattle86, NE20, CW59 |
| *tp0488* | 1* | 524108 | 524906 | 798 | 54 | TPE/TEN → Mexico A, W86 |
| *tp0515* | 1 | 557164 | 559063 | 1899 | 24 | TEN/TPE → Nichols-lineage clade |
| *tp0548* | 1* | 596164 | 596828 | 664 | 57 | TEN/TPE → Nichols-lineage clade, W86 |
| *tp0558* | 1 | 607533 | 607953 | 420 | 4 | TEN/TPE → SS14-lineage clade and syphilis ancient genomes |
| *tp0621* | 1 | 676649 | 676897 | 248 | 137 | TEN/TPE → Seattle 81-4 |
|  | 2 | 677030 | 677215 | 185 | 125 | TEN/TPE → Seattle 81-4 |
| *tp0859* | 1 | 939420 | 939442 | 22 | 11 | External sources → TPE |
| *tp0865* | 1* | 946548 | 946950 | 402 | 19 | TEN/TPE → CW86, Sea86, NE20, CW59 and Seattle 81-4 |
|  | 2* | 947238 | 947556 | 318 | 26 | TEN → CW86, Sea86, NE20, CW59 and Seattle 81-4 |
| *tp0896* | 1* | 977297 | 977302 | 5 | 4 | All primate strains → NL14, CW59 |
| *tp0967* | 1 | 1052520 | 1053803 | 1283 | 22 | TEN/TPE → Seattle 81-4 |
| *tp0968* | 1 | 1053915 | 1055118 | 1203 | 33 | TEN → Seattle 81-4 |
| *tp1031* | 1 | 1128223 | 1128350 | 127 | 19 | External sources → SS14-lineage clade |

We detected 17 genes with one recombinant region and 3 genes with more than one: *tp0136* with 7 regions, *tp0621* and *tp0865* with 2 recombinant regions each (Table 2). Interestingly, the sequences corresponding to the putative recombinant regions detected in two genes, *tp0859* and *tp1031* (Table 1), could not be found in any public database, and therefore most probably resulted from an external horizontal gene transfer event from a so far unidentified *Treponema* subspecies. Strikingly, the most likely scenario for the majority of the observed recombinant genes suggests an inter-subspecies transfer from TPE/TEN to TPA. However, one recombinant region in the *tp0136* gene corresponded to an intra-subspecies transfer within TPA (Table 1).

The inclusion of ancient genomes in our study allowed us to explore the role of these strains in recombination and their impact on current patterns of *T. pallidum*’s genetic diversity. Overall, we found eight recombinant regions and events involving the ancient genome lineages, four of these involving W86, the newly sequenced ancient genome. One of these genes, *tp0488*, was found in previous studies to have an unusual sequence in the strain Mexico A (40,49,50), which clusters with TPA sequences; this sequence was identical to TPE/TEN strains. Our study indicates that this region in *tp0488* was most probably transferred from TPE/TEN to both Mexico A and W86 lineages. An event in the *tp0179* gene was detected with the W86 and PD28 strains and the modern Nichols lineage clade as recipients and the TPE/TEN clades as putative donors. Furthermore, the W86 lineage was involved in two additional recombination events detected in the *tp0548* and *tp0558* genes. For the event involving *tp0548*, the W86 lineage and Nichols lineage clade were the recipients whereas TPE/TEN were the putative donors. For the event in *tp0558,* all TPA ancient genomes and the common ancestor of the complete SS14 lineage clade were recipients and TPE/TEN the putative donors. The lineages of the ancient genomes 94A and 94B strains were also involved in the aforementioned event detected in *tp01031,* whereas all the other strains from the SS14 lineage were the putative recipients from an external transfer. In the events detected for the genes *tp0179*, *tp0488*, *tp0548* and *tp1031*, the other TPA ancient genomes might also have been involved, but the missing data for the SNPs that define the recombination event precludes a stronger inference.

Additionally, we compared the recombination analyses with the four SRB genome datasets generated (see supplementary Note 4) and the previously described results obtained with the MRB genome alignment (Table 3). The number of SNPs detected per gene and the results of the topology tests conducted with PIM are detailed in the Supplementary Tables 7-8.

**Table 3.** Summary of recombination results obtained by PIM for the MRB alignment and the diverse SRB genome datasets

| Dataset | Number of genes | Number of genes with > 3 SNPs | Number of genes analyzed by PIM | Number of genes with phylogenetic signal | Number of genes with reciprocal incongruence | Number of genes detected as recombinant |
| --- | --- | --- | --- | --- | --- | --- |
| MRB | 1161 | 317 | 317 | 160 | 91 | 20 |
| CDC2-SRB | 1125 | 302 | 230 | - | 47 | 10 |
| BosniaA-SRB | 1122 | 301 | 229 | - | 46 | 11 |
| Nichols-SRB | 1035 | 306 | 238 | - | 48 | 12 |
| SS14-SRB | 1032 | 289 | 226 | - | 43 | 11 |

The first PIM step, which evaluates phylogenetic signal, could not be performed on the four SRB genome datasets due to missing data. Genomes distant from the reference genome often lacked reads mapping to several genes present in the references. Consequently, the analysis proceeded directly with the assessment of reciprocal incongruence in genes containing more than 3 SNPs (Table 3). This approach identified between 10 and 12 recombinant genes per dataset (Table 3). All these genes were also detected as recombinant in the MRB genome alignment, with no additional recombinant genes found beyond those identified using the MRB genome alignment. This indicates that the MRB genome alignment did not produce any false negatives while detecting nearly twice as many putative recombinant genes.

### Phylogenetic reconstruction

To build a vertical-inheritance genome phylogeny, we removed the 20 recombinant genes detected in the MRB genome alignment, as well as three genes that are hypervariable and/or subjected to gene conversion (*tp0316*, *tp0317,* and *tp0897)*, from the original alignment (1,141,136 bp with 4,822 SNPs, Supplementary File 5). The resulting alignment encompassed 1,106,409 bp with 3,047 SNPs (Supplementary File 6). Both multiple genome alignments were used to construct maximum-likelihood trees. The removal of non-vertically inherited genes had a notable effect on the phylogenetic reconstruction of *T. pallidum.* The topologies of the two ML trees, with and without these loci, are compared in Figure 2.

**Figure 2.**
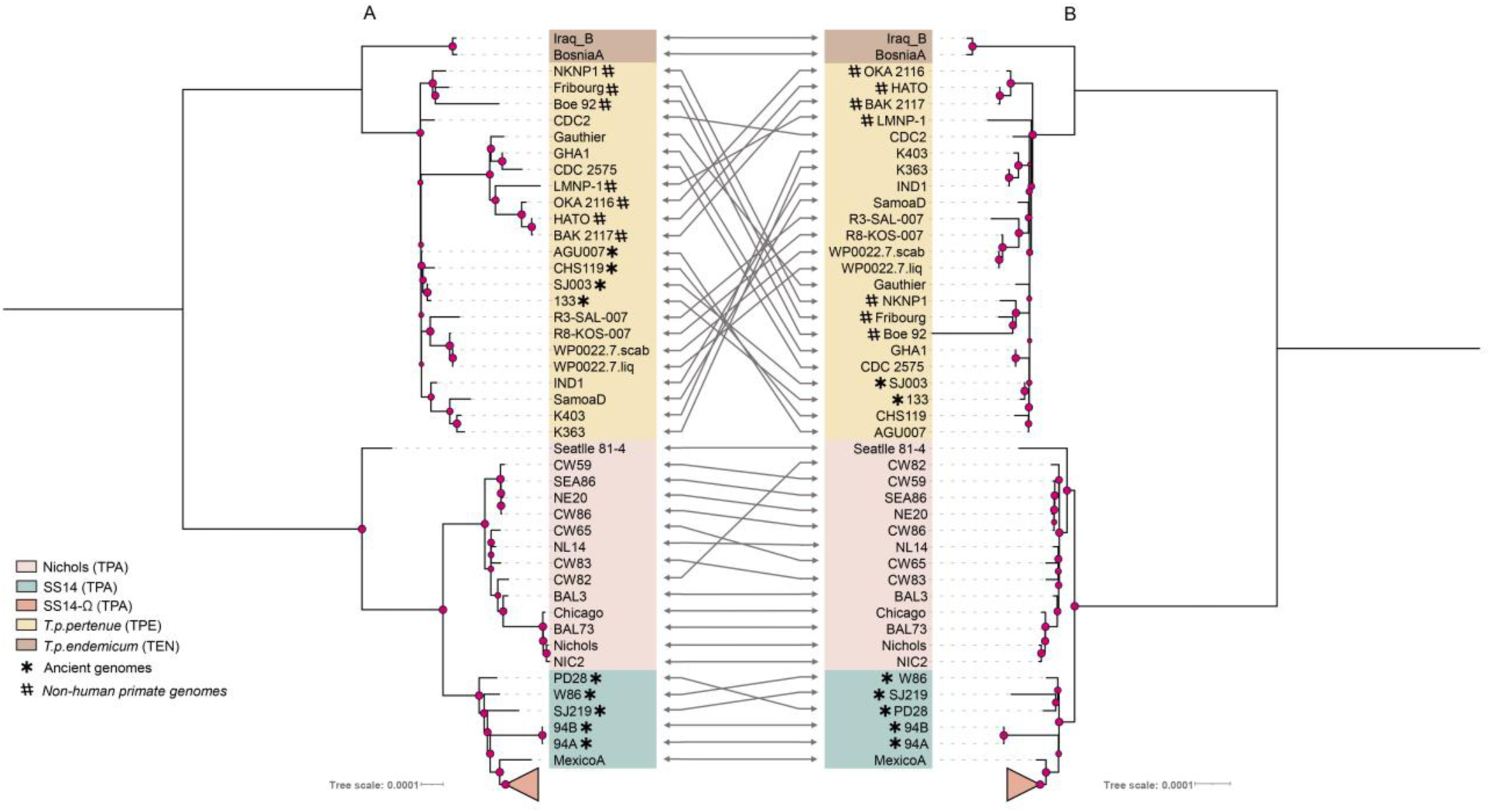
Comparison of topologies between the two maximum likelihood trees from the MRB genome alignment, A) obtained with all genes included in the whole-genome alignment, and B) obtained after excluding *tp0897, tp0316, tp0317,* and 20 recombinant genes from the whole-genome alignment. The different clades corresponding to yaws (TPE) and bejel (TEN) subspecies, and the Nichols and the different SS14 lineages of the syphilis clade (TPA) are indicated with colors, according to the corresponding color legend. Bootstrap support values higher than 70% are indicated by red circles, which are larger in better supported nodes. To enhance the clarity of the ML tree visualization, the clade comprising clinical and modern SS14-Ω strains, which exhibits minimal variation compared to other *T. pallidum* clades, has been collapsed (to see the full ML tree, see Supplementary Figure 6).

In both phylogenetic trees, a consistent classification of strains into the three subspecies and the two main lineages of TPA (Nichols and SS14) is evident. However, a noteworthy deviation occurs in the TPE clade, where specific strains (Gauthier, GHA1, CDC-2575, LMNP-1, OKA_2116, HATO, and BAK_2117) no longer constitute a well supported subclade in the tree without recombination (Figure 2B), contrasting with their arrangement in the whole genome tree (Figure 2A).

The phylogenetic positioning of the ancient SS14 genomes (94A, 94B, SJ219, PD28, and W86) remains consistent in the reconstructed tree with all genes, placing them all within the SS14 clade (Figure 2A). However, minor adjustments are noted for SJ219, PD28, and W86, with PD28 no longer being basal to all ancient genomes (Figure 2B). Notably, after excluding recombinant genes, the new historical W86 genome maintains a robustly supported basal position in the SS14 lineage, affirming its initial classification as a TPA genome (Figure 2).

Nonetheless, several topological incongruities emerged between the two ML trees obtained (Figure 2) for the Nichols subclade. The unexpected location of Seattle 81-4 in the whole genome tree, distanced from all TPA genomes and basal to the joint Nichols and SS14 lineages, undergoes a shift after removing the genes mentioned above (Figure 2B) to occupy a basal position within the Nichols lineage. Moreover, strain CW82, originally situated within it in the ML tree obtained using the whole genome alignment (Figure 2A), is now firmly situated at the base of the Nichols lineage in the tree obtained using the recombination-free alignment (Figure 2B).

Interestingly, for this study we analyzed two pairs of samples: 1) IND1 and K363, and 2) Nichols, and NIC2; each pair originating from the same clinical sources but sequenced using different methods. NIC1 and Nichols were originally from the same sample which was cultured through multiple rabbit passages before sequencing (42,51). Similarly, IND1 and K363 also come from the same original sample, but while K363 underwent rabbit passage, IND1 did not (42,47). Comparative analysis revealed SNP variations between both pairs of samples, with notable discrepancies in the phylogenetic tree placements of IND1 and K363. These differences were primarily attributed to flagged recombinant genes, emphasizing the need for careful interpretation of genomic data. For further details on these analyses, see Supplementary Note 3.

### Natural selection analysis

To study the effects of natural selection, using the MRB genome alignment, from the total set of 1161 genes (Supplementary Table 3), 317 genes with three or more SNPs were analyzed in HyPhy, using aBSREL, a "branch-site" model to detect positive selection. This analysis identified 28 genes showing evidence of positive selection (Supplementary Table 9 and Figure 3), all of which have a large number of SNPs in the strains comprising our dataset. These genes included 10 putatively recombinant genes (*tp0131, tp0136, tp0462, tp0621, tp0856 tp0865, tp0859, tp0967, tp0968* and *tp1031*). Although most branches in which these recombinant genes were found to be under positive selection corresponded to the deeper branches of the phylogeny, including the branch leading to the MRCA of the TPE/TEN subspecies, in some cases the signature of selection was limited only to branches leading to the specific strains found to be involved in recombination events of these genes.

**Figure 3.**
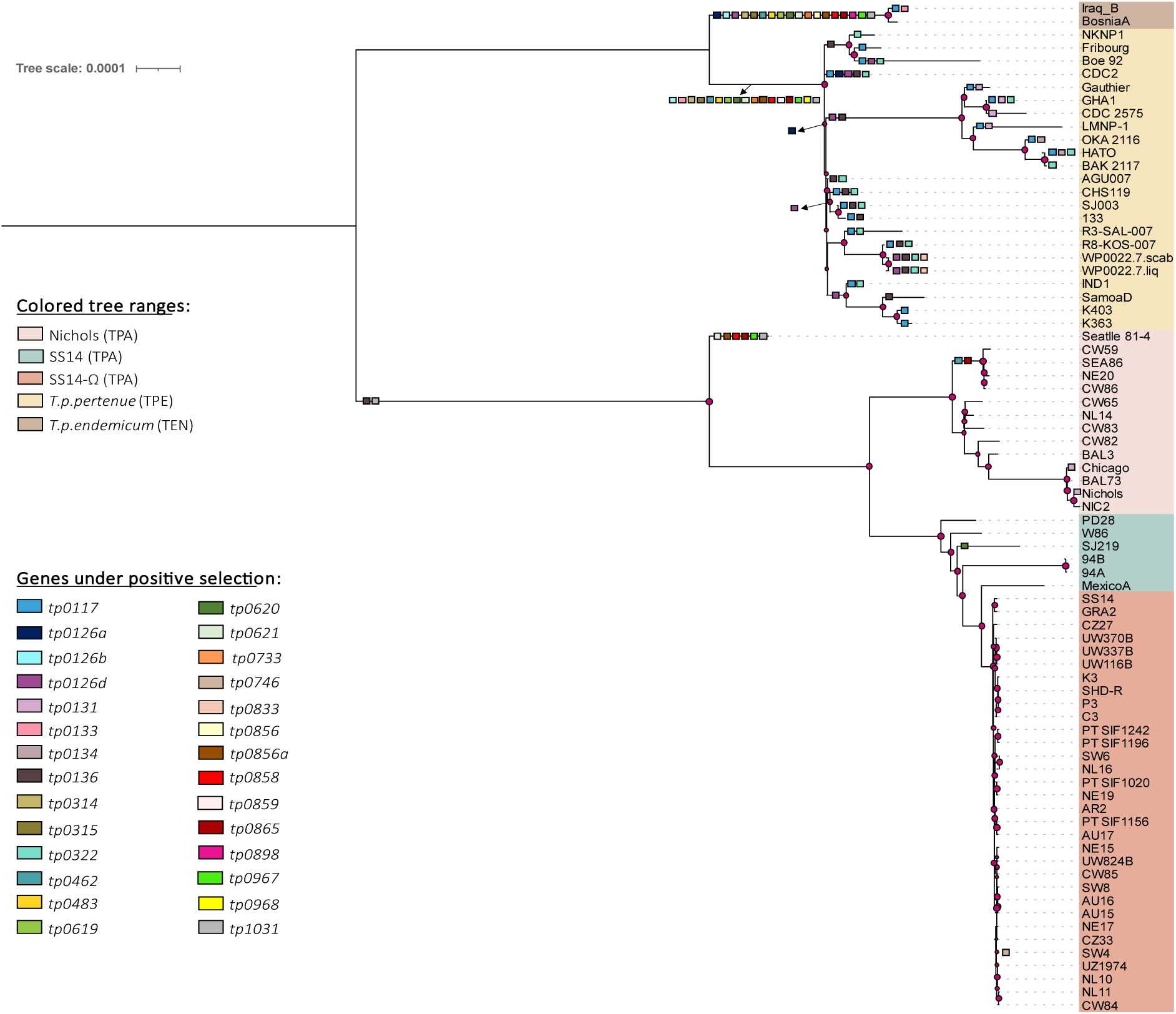
Maximum likelihood tree derived from the MRB genome alignment, obtained with all genes included in the whole-genome alignment, in which genes found under positive selection have been represented on their branches. For more details about the analysis results see Supplementary Table 9.

Interestingly, despite poor coverage of the *tpr* genes in some genomes, apart from two *tpr* genes that had also been detected previously as recombinant (*tp0131 and tp1031)*, we found two more genes of this family (*tp0620* and *tp0117)* with evidence of positive selection. For the locus *tp0620* (*tprI)*, the branch leading to the MRCA of the TPE and TEN strains in the phylogeny appears to be under positive selection. This gene has been previously described (47) as having a modular genetic structure in certain *T. pallidum* strains. This structure differs from that of other *T. pallidum* subspecies, which might explain the detection of positive selection. Moreover, aBSREL detected the *tp0117 (tprC)* gene to be under positive selection in the branch leading to the MRCA of TPE and TEN strains, and for the recombinant *tp0131 (tprD)* gene in the branch leading to the MRCA of Nichols, Chicago, LMNP-1, Gauthier, and CDC_2575 strains. Previous studies have considered these two genes as paralogs (47,49), created by a gene conversion mechanism that would have copied a portion of the *tp0117* gene into *tp0131*, thus explaining differences between these genes in some strains (Gauthier, LMNP-1, CDC_2575, Nichols and Chicago).

We also detected evidence of positive selection in loci *tp0314* and *tp0619* in the branch leading to the MRCA of TPE and TEN strains. This could be a consequence of a previously described gene duplication (52), in which a paralogous sequence covering the *tp0314* and *tp0619* genes was found to be almost identical to the region containing the two *tpr* genes *tp0620*-*tp0621* (*tpr*IJ) in all the TPE and TEN strains analyzed.

In addition, other loci (*tp0856*, *tp0856a,* and *tp0858*) also showed a modular structure in previous analyses (47), apart from the putative recombinant genes *tp0136* and *tp0620* (*tprI*), both previously described. All of these genes were found to be under positive selection in our analysis. The modular nature of these genes and the branches detected to be under positive selection together suggest that the recombination events occurred through gene duplication and gene conversion within treponemal genomes, and that they could result in substantial changes in gene and protein sequences.

Furthermore, 11 non-recombinant genes were detected to be under positive selection in the branch leading to the MRCA of TPE and TEN strains. Two other non-recombinant genes, *tp0134* and *tp0833,* were detected to evolve under positive selection in the branch leading to the MRCA of the historical genomes of TEN. Specifically, only the branches leading to the OKA2116 and HATO strains for the *tp0134* gene and the branches leading to the WP00227.liq and WP00227.scab strains for the *tp0833* gene exhibited signal of positive selection.

As we observed a close relationship between recombination and selection, we also examined the functional roles of the proteins coded for by the recombinant and additional genes found to be under positive selection, according to Uniprot and a literature search (detailed in Supplementary Table 10). Despite some proteins having an unknown function, most of the proteins identified here appear to play an important role in the defense of the pathogen against the host immune system and are potentially involved in virulence.

### Molecular clock dating analysis

Molecular clock dating analysis (Figure 4) using an uncorrelated lognormal relaxed clock and a Bayesian skyline population model was performed on the dataset of 75 genomes (sequences IND1 and NIC2 were removed as duplicates to avoid overinflation of genetic signal from these samples). We also removed the 23 recombinant or hypervariable genes as described above. The alignment was reduced to only variable sites (3,047 bp) to facilitate handling and computation. The inferred time to the most recent common ancestor and the 95% Highest Posterior Density (HPD) intervals estimated for the major *T. pallidum* clades are detailed in Table 3.

**Figure 4.**
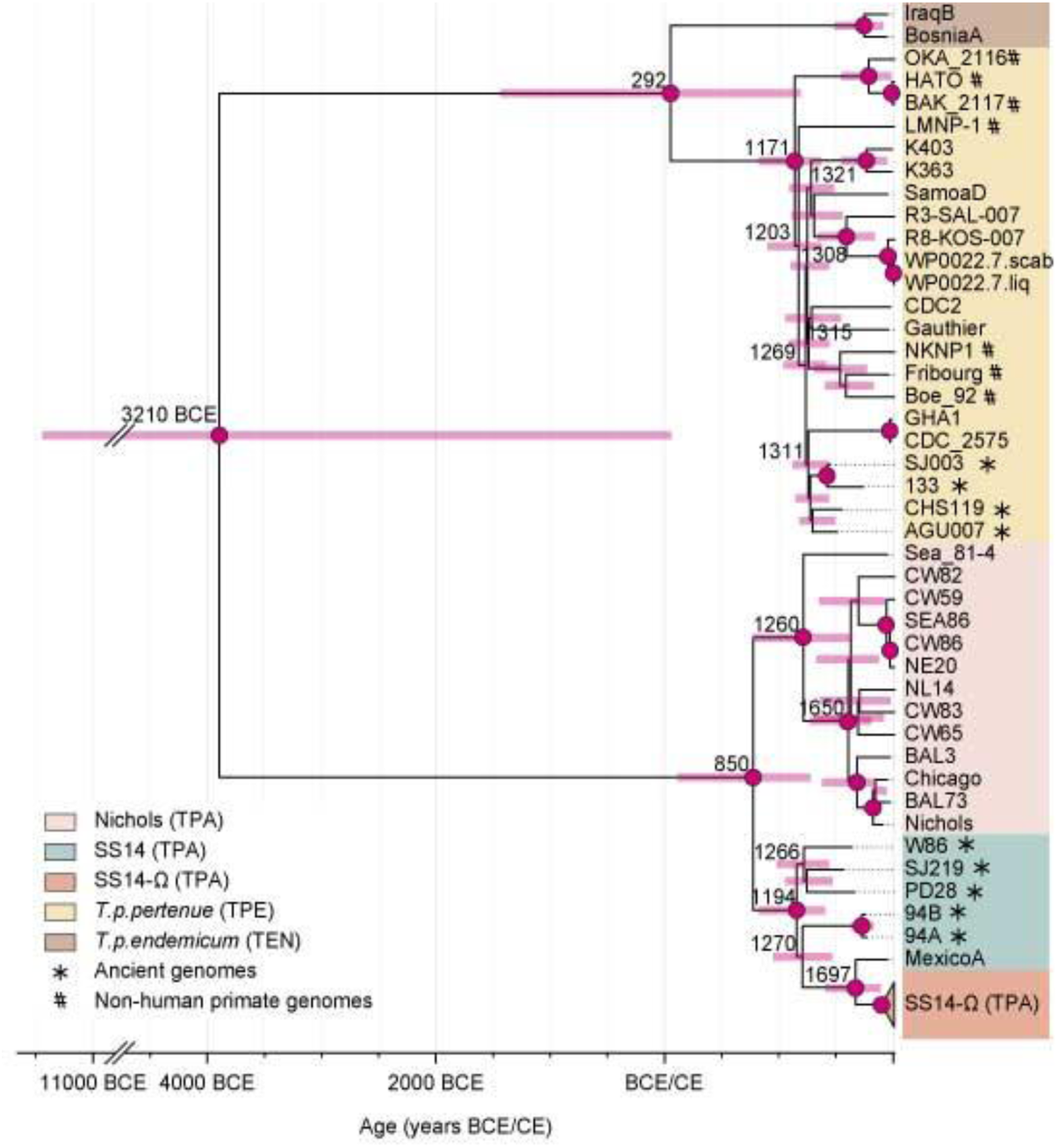
Maximum Clade Credibility tree representing the time-aware Bayesian phylogeny of *T. pallidum* estimated using BEAST 2. See main text for further details on the data employed in this analysis. Numbers denote the median age estimates for the main nodes. Pink bars show the 95% HPD of the node age estimate. The different clades corresponding to the yaws (TPE) and bejel (TEN) subspecies, and the Nichols and the different SS14 lineages of the syphilis clade (TPA) are indicated in the figure with colors, according to the corresponding color legend. Bayesian posterior probabilities higher than 95% are indicated by red circles. The clade comprising clinical and modern SS14-Ω strains was collapsed to improve readability (see Supplementary Figure 7 for the fully dated tree).

The divergence time between the TPE/TEN and TPA, i.e. the tMRCA for the entire *T. pallidum* family, was broadly estimated to fall between 9430 BCE and 60 CE (median 3210 BCE), in accordance with a previous estimate (53). Consistent with previous studies (39,53), we dated the tMRCA of all TPA lineages between 120 CE and 1280 CE (median 850 CE), the TPA Nichols lineage between 1270 and 1810 CE (median 1650 CE), and between 780 and 1650 CE (median 1260 CE) when also including Seattle 81-4 as part of the Nichols lineage. The SS14 lineage was dated to between 830 and 1410 CE (median 1190 CE) and lineage SS14-Ω to 1820 - 1965 CE (median 1910 CE). The TPE/TEN MRCA was estimated between 1430 BCE and 1200 CE (median 290 CE) and the MRCA of all TPE genomes between 830 and 1380 CE (median 1170 CE). All these nodes obtained a Bayesian posterior probability of at least 0.99.

The median evolutionary rate was estimated to 4.56×10^-5^ substitutions/SNPsite/year (95% HPD interval: 2.84 - 6.53×10^-5^), which corresponds to 1.26×10^-7^ substitutions/site/year (95% HPD interval: 0.78 - 1.8×10^-7^) for the non-SNP-restricted *Treponema* genomes with hypervariable and recombining genes excluded.

## DISCUSSION

### The relevance of the right reference genome

In this study, we examined the impact of genomic reference selection on the phylogenetic and evolutionary analyses of *T. pallidum*. Our findings highlight the importance of this choice, as using a inappropìate reference genome for mapping HTS reads can introduce errors that affect downstream analyses, such as recombination detection and phylogenetic inference. This is consistent with previous research by Valiente-Mullor *et al*. (37) which showed that relying on a single reference in microbial genomics can lead to inaccuracies, especially when dealing with genetically diverse isolates.

To date, no previous study has conducted a comprehensive evaluation of the impact of utilizing various genomic references in the mapping process to obtain complete genomes of *T. pallidum*. Specifically, there has been little focus on how this affects the integration of ancient genomes with modern genomes of this bacterium. In the study by Pla-Díaz *et al*. (54), the influence of three distinct references (CDC2, Nichols, and SS14) on recombination detection was assessed with a dataset encompassing clinical and modern genomes of the three *T. pallidum* subspecies. The conclusion drawn was that recombination detection remained consistent irrespective of the genomic reference chosen. However, BosniaA was not included in that investigation as an additional potential genomic reference because it was the only genome available representing the TEN subspecies at that time. Furthermore, the genomic diversity, measured in terms of variation, was lower in the SRB genome dataset used by Pla-Díaz *et al*. (2,625 SNPs) compared to the 4,822 SNPs present in the final MRB genome alignment utilized in the current study. Moreover, our updated genome database includes not only modern genomes, as in Pla-Díaz *et al*., but also ancient genomes.

Our results further indicate that the choice of genomic reference could substantially affect the reconstruction of ancient genomes, thereby influencing the resulting phylogenetic inferences. Specifically, failure to select the closest-reference genome for *T. pallidum* may significantly impact phylogenetic reconstruction, particularly when applied to a dataset containing genomes from all three *T. pallidum* subspecies, and when including ancient genomes (Supplementary Figures 2-5). While the calculation of average RF distances in pairwise tree comparisons reveals relatively similar means, a closer examination of the ML trees derived from each of the four SRB genome datasets against the ML tree obtained using the MRB genome alignment, reveals significant topological incongruities. This is particularly evident in the placement of ancient TPA genomes and the Mexico A strain.

The Mexico A strain genome, derived from rabbit cultivation and dating back to a sample taken in 1953, is closer to the ancient genomes which generally fall basal to the SS14 strains, than to the SS14-Ω sublineage that includes the majority of the contemporary clinical strains of the clade. Unlike the clonal SS14-Ω strains, ancient TPA genomes and Mexico A exhibit greater genomic variation, affecting consensus genomic sequences when using different references. In the case of Mexico A this variation might result from microevolution during rabbit culturing, as shown in a 2023 study by Edmondson et al. (29). Consequently, many studies exclude rabbit-cultivated strains from molecular clock dating to avoid phylogenetic noise. However, two new genomes (MD18Be and MD06B), not cultivated in rabbits from clinical strains isolated in the USA in 1998, cluster with Mexico A (55), suggesting that much of Mexico A’s variation is genuine. Due to time constraints, some of the most recently published genomes were omitted from our analyses. These genomes deserve further exploration and should be incorporated in future studies on the impact of closely related genome references, especially in the context of ancient *T. pallidum* genome analyses.

In this study, we also aimed to investigate the potential impact of using different genomic references on recombination detection. We compared genes identified as recombinant using the PIM method across four SRB genome datasets reconstructed with distinct *T. pallidum* genomic references (CDC2, BosniaA, Nichols, and SS14) against recombination detected in the MRB genome alignment generated by our new mapping strategy utilizing the closest genome reference. The findings revealed that none of the SRB genome datasets identified new recombinant genes not detected in the MRB genome alignment. However, many of the 20 genes identified with PIM in the new dataset generated by the revised mapping strategy were not found in the four SRB genome datasets due to substantial missing data in the gene sequences. This underscores the critical importance of selecting a closely related genomic reference for achieving accurate and high-quality genomic sequences. Using a closely related reference genome not only enhances the accuracy of the reconstructed sequences but also significantly improves the overall genome coverage.

As we delve into the importance of genomic references for mapping accuracy, evaluating the reliability of sequencing techniques is also important. For this purpose, we included in our genome dataset four selected samples—IND1 and K363 from one clinical specimen, and Nichols and NIC2 from another—that were obtained in previous studies (42,47) using different sequencing and/or processing methods. Based on the obtained results (see Supplementary Note 3), notable differences are evident among the sequences derived from strains originally sourced from the same sample, particularly noticeable between strains IND1 and K363. This underscores the critical importance of obtaining high-quality sequences and highlights how errors in sequencing techniques, as well as DNA enrichment through rabbit passage, can significantly impact subsequent sample analyses.

Given the absence of a standardized culture method for *T. pallidum*, Whole Genome Amplification (WGA) is a PCR-based technique that is effective for modern samples, especially when dealing with low quality DNA. However, as demonstrated by Forst *et al.* (56), WGA can struggle with the low DNA concentrations typically found in ancient DNA (aDNA) extracts. This limitation is similar to the challenges faced by PCR in general. Therefore, WGA is not recommended for ancient samples. In contrast, in-solution target enrichment is the most effective method for ancient genomes and low-quality samples, yielding consistent results across studies (12,19,24–27,39,46). However, while this method is beneficial for ancient samples, it may introduce bias if the probes are designed using a reference genome that differs significantly from the sample being enriched, potentially leading to a loss of authentic DNA fragments. This issue was observed in the IND1 sample, a TPE sample for which TPA-specific baits were used (42). The choice of enrichment technique is crucial, depending on the sample’s characteristics, to avoid the potential loss of authentic *Treponema* DNA fragments or other biases.

### Insights into *T. pallidum* evolution

Despite the persistent challenges in elucidating recombination mechanisms in *T. pallidum*, various studies (47,49,52,54,57–62) have explored the occurrence of recombination in this bacterium and the potential impact of natural selection on the transferred genetic material. Using the PIM method (27,42,53,63–67), we identified 28 recombinant regions in 20 different genes, unveiling a new recombinant gene *(tp0621)* not previously detected with this method (25,27,42,54,63). Despite revealing this new recombinant gene, the enhanced quality of the draft genomes achieved through our novel mapping approach only partially addresses the considerable number of missing positions in certain loci, which for modern samples stems from the unavailability of a standard culture system, and for the ancient samples from the highly degraded nature of ancient DNA. For both, mapping short reads is particularly challenging for paralogous and repetitive genomic regions, as observed in some *tpr* genes and underscores the complexities associated with working with low-quality genomes.

The observation that all identified recombination events involve transfers between subspecies (TPE/TEN to TPA), except for a notable occurrence in the *tp0136* gene between the Nichols and SS14 TPA lineages, aligns with the results in Pla-Díaz *et al*. (54). This is compelling due to the specific geographical niches TPE and TEN occupy today. To consider the possibility of recombination between TPE/TEN and TPA strains, we must assume the coexistence of diversified clades in a sympatric manner, allowing them to simultaneously infect a common host. Yet, there is currently no evidence of human coinfection. Furthermore, the involvement of ancient genomes from both TPE and TPA lineages in recombination events across eight different genes adds complexity to the puzzle. Key questions that emerge are the locations and timelines of these intriguing events.

Recent indirect evidence from ancient genomes suggests a prolonged and intricate coexistence of syphilis and yaws in historical Europe (24,68,69). Despite this, the origin and divergence of *T. pallidum* subspecies remain debated among scientists and historians. Using an enhanced dating approach, we estimate the time to the most recent common ancestor (tMRCA) for the entire *T. pallidum* family to be between 9430 BCE and 60 CE, with a median of 3210 BCE. Additionally, a recent study by Majander *et al*. (27) confirms the presence of the TEN subspecies of *T. pallidum* in the Americas before Columbus’ arrival. However, there is still no evidence for the presence of TPA or TPE in the Americas prior to this event, leaving the origins of these subspecies unresolved.

Based on our divergence time estimates—850 CE for TPA and 290 CE for TPE/TEN—and detected recombination events between subspecies, there are indications of potential geographic overlap facilitating genetic exchange, although direct evidence is lacking. The diversity and wide geographical span of contemporary lineages in early modern Europe reduce the likelihood of yaws and syphilis being newly introduced simultaneously. Some exchanged genes may have facilitated rapid adaptation to environmental or host behavioral changes, as revealed by the natural selection analysis. Many recombinant genes have undergone evolution through positive selection, with most playing crucial roles in virulence and immune evasion (see Supplementary Table 10). However, the exact functional roles of these gene variants in transmission and virulence remain unclear, limiting the explanatory power of ancient genetic variation.

While our natural selection analysis follows standard approaches to test for positive selection, we acknowledge that the presence of recombination within genes could potentially confound these results. A single phylogenetic tree may not fully capture the evolutionary history of recombinant genes, leading to possible biases in branch-length estimation and, consequently, in the detection of positive selection. To address this problem, we took this into account by using the phylogenetic tree obtained for each gene in the inference of positive selection, following Pla-Díaz *et al.* (54). A more detailed analysis of natural selection throughout the *T. pallidum* genome, beyond the scope of this work, will benefit from independently analyzing the recombinant and non-recombinant regions of the recombinant genes, as we did in previous work (54). Considering the confirmed presence of TPE and TPA in early modern Europe as distinct species at the genomic level, genetic exchange between subspecies likely facilitated the global spread and adaptation of these diseases. However, these hypotheses require further substantiation with robust evidence.

## CONCLUSIONS

Our study thoroughly examines the impact of choosing an appropriate genomic reference on the phylogenetic and evolutionary analyses of *T. pallidum*. We observed that while recombination detection was consistent across different references, the selection of a specific reference had a substantial effect on the reconstruction of ancient genomes and the resulting phylogenetic interpretations. These findings suggest that creating an artificial most recent common genome, akin to what has been done for the *M. tuberculosis* complex (70,71), could be advantageous for future mapping efforts. Such a development could help reduce biases from mapping strategies and improve the accuracy of ancient *T. pallidum* genomes. Additionally, we identified new recombination events, positive selection targets, and refined dating estimates for significant events in the species’ history. Our work underlines the importance of recognizing methodological implications and reference genome bias in HTS-based whole-genome analysis, contributing to a more profound understanding of *T. pallidum* and its subspecies.

## MATERIALS AND METHODS

A summary of the workflow used in the analysis of the 77 *T. pallidum* genomes is shown in Figure 5.

**Figure 5.**
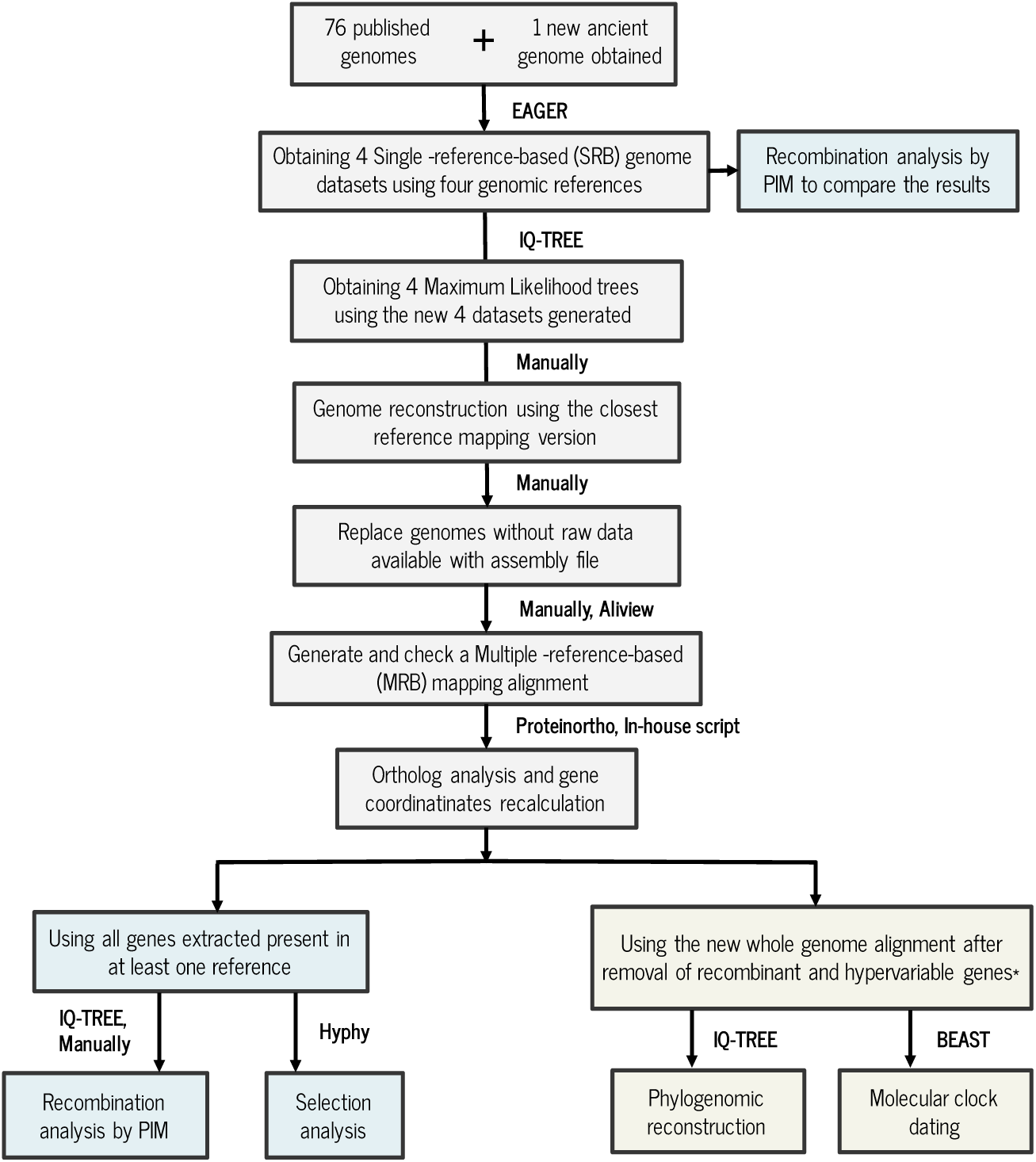
Analysis workflow for the genomic and phylogenomic analysis of the 76 previously published *T. pallidum* genomes and one new historical genome obtained in this study (W86). The hypervariable genes indicated by * are *tp0897* and *tp0316*.

### Sample processing

The upper-left premolar tooth sample was collected from human remains at the Wrocław University of Environmental and Life Sciences archaeological collection as part of the study project focusing on Wrocław’s 17th century population genetics. Treponemal infection was not identified anthropologically. The sample was pretreated in clean room facilities, dedicated to state-of-the-art ancient DNA work, at the Museum and Institute of Zoology, Polish Academy of Sciences, in Warsaw. To avoid possible human and environmental DNA contamination, the surface of the tooth was sanded off with a hand rotary tool. The tooth was then washed with 5% sodium hypochlorite, molecular-grade water and 75% ethanol, UV-irradiated for 10 minutes on each side and then pulverized using a Retsch MM200 mixer mill. All sampling tools and all reusable items were regularly cleaned with diluted bleach and UV irradiated between each use to minimize the risk of contamination. As a part of the project, screening for specific pathogens was conducted using a PCR-based test for selected genetic markers (See Supplementary Note 1).

### DNA extraction and library preparation

DNA extraction (72) and double-stranded Illumina library construction were performed according to an established protocol (73,74) in the clean room facilities dedicated to ancient DNA processing at the Institute of Evolutionary Medicine in Zurich (75). Library pools were shotgun sequenced with an Illumina NextSeq platform using a NestSeq 500 Mid Output Kit (75 cycles paired-end) for the initial pathogen screening.

### UDG treated libraries

Subsequently, to remove ancient-DNA-specific DNA damage, the Uracil-DNA glycosylase (UDG) (76) was used to treat the additional libraries prepared from the same DNA extract.

### Whole genome capture

A custom target enrichment kit (Arbor Biosciences) was used for the whole genome capture as in Majander *et al.* (25). For this purpose, 60 bp long RNA baits with a 4 bp tiling density and 99% identity were designed based on a selection of representative genomes (Nichols: GenBank CP004010.2, Fribourg Blanc: GenBank CP003902, SS14: GenBank CP000805.1) from each *Treponema pallidum* subspecies or clade. UDG-treated libraries were pooled in equimolar concentration, and 500 ng final pools were hybridized in 60°C for 48 hours following the manufacturer’s instructions. 10 nM capture pools were sequenced using an Illumina NextSeq 500 High-throughput Kit (75 bp paired-end). Sequencing reads obtained for each UDG-treated, enriched library were processed with the EAGER pipeline (v.1.92.55) (36). After removing the adapter sequence using AdapterRemoval v. 2.2.1a (77), libraries from the W86 individual were merged and processed as paired-end sequencing reads. The authenticity of ancient DNA was assessed by EAGER analyzing C to T deamination at the terminal base of the DNA fragment.

### Dataset selection and read processing

We generated a genomic dataset comprising 68 modern *T. pallidum* draft genomes (47 TPA, 19 TPE and 2 TEN) from previously published studies, in addition to 8 published historical genomes, and one new ancient genome we obtained (W86). The modern genomes were selected as a representative set of the diversity patterns observed in the phylogenetic trees reconstructed at the start of this study in 2020-2021. The ancient draft genomes selected were those with a minimum coverage of 5X among those published up to that date (24–26,78). Out of the total of the 76 genomes selected for analysis, raw sequencing data were available for 61, while for the other 15 only the consensus genome sequences files could be obtained (see Supplementary Table 1). The raw data and the consensus genome sequences were downloaded from the NCBI and ENA databases (79,80). For the 15 genomes without raw data available, HTS-like reads were simulated based on their genome assembly files using the tool Genome2Reads (integrated in the EAGER pipeline (36)) for a posterior comparison with their assembly consensus genome sequences available and downloaded from the public databases.

To reconstruct all the individual genomes from the raw short-read data (including the simulated raw data for 15 samples without available raw data) we carried out raw read quality control and preprocessing, removed duplicates and identified variants using the programs implemented in the EAGER pipeline (v.1.92.55) (36), as in previous studies (25,36,42). After processing the de-multiplexed sequencing reads, sample sequencing quality was analyzed with FastQC version 0.11.5 (81). Following processing by AdapterRemoval ver. 2.2.1a (77), the mapping was carried out using CircularMapper version 1.0 (36), with default BWA (-l 32, -n 0.04, q 37) parameters (82) and Nichols (NC_021490.2) and SS14 (NC_010741.1) genomes, which represent the two main groups of TPA, and the CDC2 (NC_016848.1) and BosniaA (NZ_CP007548.1) genomes, which are well-studied TPE and TEN strains, respectively, as reference. Each of the genomes in the dataset was mapped to each of these four references. The *MarkDuplicates* method provided by the Picard Toolkit (2019) was applied to remove duplicate reads and DamageProfiler version v0.3.12 was utilized to estimate the DNA damage parameters for the new ancient genome obtained (83). Indel realignments were performed using GATK (version 3.6) (84) and single nucleotide polymorphisms (SNPs) for the resulting mappings were called using GATK UnifiedGenotyper with the following parameters for SNP calling: -nt 30, -stand_call_conf 30, --sample-ploidy 2, -dcov 250, --output_mode EMIT_ALL_SITES; and the following parameters for SNP filtering: DP>5, QUAL>30. The reconstructed W86 genome and its main features were represented graphically using BRIG (85).

### Antibiotic resistance

Two mutations on the 23S ribosomal RNA operon, A2058G and A2059G (46,86), were investigated to assess macrolide azithromycin resistance in the newly obtained ancient genome W86. For this purpose, 23S rRNA gene sequences for operons 1 and 2 with 200 bp added to the 5’ and 3’ flanking regions were extracted from the Nichols reference genome and aligned to the W86 genome. Subsequently, the presence or absence of each of the two mutations was assessed with variant calling.

### Obtaining the different Single-reference-based (SRB) genome datasets

To assess the influence of different genomic references on mapping in terms of phylogenetic assignment, we conducted the following analysis. We used the 77 samples selected for this study to create four distinct SRB mapping-genome datasets. Each sequence in these datasets was mapped using each of the four different genomic references, corresponding to the three *T. pallidum* subspecies and the Nichols and SS14 clades: Nichols, CP004010.2; SS14, NC_021508.1; CDC2, CP002375.1 and BosniaA, CP007548.1. For the 15 genomes without raw data available, HTS-like reads were simulated based on their genome assembly files using the tool Genome2Reads (integrated in the EAGER pipeline (36)). The parameters used are consistent with those outlined in the Dataset Selection and Read Processing section. Henceforth, we will denote each of these SRB genome datasets as Nichols-SRB, SS14-SRB, CDC2-SRB, and BosniaA-SRB dataset, respectively.

For each of the 4 generated SRB genome datasets, a Maximum likelihood (ML) tree was generated with IQ-TREE (version 1.6.10) (87), using GTR+I as the evolutionary model with 1000 bootstrap replicates. Regarding the newly acquired ancient genome, W86, its position in each of the phylogenetic trees derived from the four SRB datasets was examined to ascertain its classification within the three subspecies and/or clades of *T. pallidum*.

### Multiple-reference-based (MRB) final genome alignment

As it is known that SNP calling in a genome is dependent on the choice of the reference used for mapping (37), we carried out a proximity evaluation to determine the closest reference for each of the genomes to reconstruct the final genome alignment to be used in the subsequent evolutionary analyses.

The process involved selecting the genomic sequence for each of the 62 strains with available raw data from the four SRB genome datasets obtained earlier. This selection was based on the strain classification into the three different *T. pallidum* subspecies and/or subclades (TPE, TEN, Nichols, and SS14) established in previous studies, which provided the genomes for this investigation. Subsequently, the chosen genome sequences for the 62 strains, according to their closest genome reference from the four genome-mapping datasets, were consolidated into a single MRB-mapping multiple genome alignment.

Next, we incorporated 15 previously assembled genomes, which were downloaded from public databases without accompanying raw data, into the single MRB-mapping multiple genome alignment we previously obtained. We chose not to use their consensus genome sequences obtained by simulating their reads in this new alignment. Instead, we believe that the assembled genomes from public databases, as obtained in their respective studies, are of higher quality than those generated by simulating reads. Our goal is to compare *a posteriori*, their placement in a phylogeny using the simulated reads versus their original assembly consensus sequences.

We realigned all sequences, producing a new whole-genome alignment comprising 76 *T. pallidum* genomes. The selected sequence mapping version for the W86 genome from the SRB genome datasets according to its closest genome reference was then added to this whole-genome alignment, which underwent realignment again. The result was a final whole-genome alignment of 77 sequences. Subsequently, using this MRB-based multiple genome alignment, we generated a maximum likelihood (ML) tree with IQ-TREE (version 1.6.10) (87), employing GTR+G+I as the evolutionary model and performing 1000 bootstrap repetitions.

Then, the four ML trees obtained before from the SRB genome datasets were compared with the whole-genome phylogeny of the final MRB alignment generated to analyze the topological differences between them.

Furthermore, to quantify the topological differences between the four phylogenetic trees derived from the SRB-mapping datasets and the whole-genome phylogeny of the MRB multiple genome alignment generated, we computed the Robinson-Foulds (RF) distance among them using RAxML version 1.2.0 (88). For this comparison, we computed the RF distance between all the trees. Additionally, we calculated the average of these distances to identify the most discrepant topology.

Additionally, an orthology analysis was carried out with Proteinortho (version V6.0b) (89) to identify orthologous genes in the four reference genomes employed for mapping. The genomic coordinates of each gene present in at least one of the four reference genomes were then calculated according to their corresponding location in the final merged whole-genome alignment.

The protein translations for all the genes present in at least one reference genome were compared to the original gff3 files of each of the four references, to ensure that the final MRB alignment generated was correct, and that no protein was accidentally truncated (Supplementary Files 7-8).

### Recombination detection: PIM

The presence of recombination in the complete genomes of *T. pallidum* could potentially interfere with the inference of phylogenetic tree topologies, as described in Pla-Díaz *et al*. (2022) (54). We thus used the PIM pipeline (54) to investigate putative recombining genes. In summary, the procedure included the following steps (with details provided in Supplementary Note 4): 1) A maximum likelihood (ML) tree was computed for the MRB genome alignment using IQ-TREE (87). 2) The 1061 genes present in at least one of the reference genomes were extracted, and the number of SNPs per gene was computed using an in-house script (Supplementary File 9), discarding genes with less than 3 SNPs. 3) For each of the genes retained, the phylogenetic signal in each gene alignment was tested using likelihood-mapping (90) in IQ-TREE, retaining only those genes that showed some phylogenetic signal (see Supplementary Note 4). 4) For the remaining genes, an ML tree was computed using IQ-TREE. Next, topology tests for each candidate gene were performed by IQ-TREE using two different methods: Shimodaira–Hasegawa (SH) (91) and Expected Likelihood Weights (ELW) (92). First, using the corresponding gene alignment, we compared the likelihood of each individual gene tree to the reference genome-wide tree (the phylogeny constructed from the multiple genome alignment). Then, we compared the same likelihoods using the entire genome alignment. When both tests reject the topology that is not derived from the corresponding alignment, this is referred to as a reciprocal incongruence (individual gene in the first comparison, the complete genome in the second). 5) For all genes passing the previous filtering steps, the presence of a minimum of three consecutive homoplasic SNPs congruent with a recombination event were checked using MEGAX (93) to classify a gene as recombinant.

Several genes have a large proportion of sites with missing data due to the challenges in mapping short read data from regions containing repetitive DNA. Most of these genes pertain to the *tpr* family, which comprises groups of paralogous genes. For each of these genes, strains with a high proportion of missing data were removed, in order to be able to still analyze these interesting genes with the PIM pipeline. The hypervariable gene *tprK (tp0897)* was completely discarded from the recombination analysis following the results in Pla-Díaz *et al*. (2022) (54) because its seven hypervariable regions undergo intrastrain gene conversion, which have been studied in detail elsewhere (94–96).

We conducted recombination analyses with the four datasets derived from each SRB-mapping dataset. The goal was to compare recombination patterns between the new mapping strategy and the use of a single genomic reference. Initially, we extracted genes from each of the four alignments, corresponding to the annotation file of the genomic reference employed for each SRB genome dataset generated. Subsequently, we applied the PIM method to assess recombination within each set of extracted genes per dataset, allowing for a comprehensive comparison of the results. It is essential to highlight that several genome sequences within each of the four SRB genome datasets exhibited a significant amount of missing data. This poses a challenge in performing the PIM recombination analysis. To address this, we carefully selected genes within each of the four SRB genome datasets, focusing on those with more than 3 SNPs. Subsequently, we proceeded directly to the topology test step without performing the likelihood mapping test.

### Phylogeny reconstruction

Prior to the final phylogenetic analyses, genes identified by PIM as recombinant were excluded from the MRB genome alignment. To avoid introducing a potential bias into the phylogenetic signal, 3 genes (*tp0897, tp0316, tp0317*) which contain repetitive regions and have previously been reported as hypervariable and/or under gene conversion (20,47,94–96) and thus induce recombination-like effects in phylogenetic inference, were also removed. As the *tp0317* gene is embedded inside the *tp0316* gene, and moreover the coordinates from the BosniaA reference genome for *tp0316* spanned a longer region compared to the other reference genomes, *tp0316* and *tp0317* were removed according to the BosniaA reference genome coordinates for *tp0316*. A maximum likelihood tree was then constructed with IQ-TREE, using GTR+G+I as the evolutionary model.

### Natural selection analysis

In addition to recombination processes, natural selection also plays an important role in genetic diversity patterns. In prior research, Pla-Díaz *et al*. (2022) (54) demonstrated a close relationship between recombination and selection in *T. pallidum*, suggesting an important role of these processes in the evolution of this bacterial species, especially in TPA lineages. Here, we tested for positive selection in a subset of 317 (out of 1161) genes that comprised three or more SNPs. If a strain in the gene alignment had more than 50% missing data, it was excluded from the analysis. Additionally, *tp0897*, *tp0316* and *tp0317* were excluded from this analysis because of hypervariable regions and the gene conversion signal present in these genes (47,49,52,94). Then, to test if positive selection occurred along the different lineages of the phylogeny (97), we employed HyPhy (version 2.5.32) (98), using the aBSREL model (adaptive Branch-Site Random Effects Likelihood), which is an improved version of the commonly-used "branch-site" models (97,99). We used default settings and the ML phylogenies of each gene to prevent the confounding effects of recombination on the inference of positive selection, as we did in Pla-Díaz et al. 2022 [54] We assessed statistical significance using a likelihood-ratio test (LRT).

### Molecular clock dating analysis

Molecular clock analysis was performed for the dataset comprising 75 genomes, because the IND1 and NIC2 sequences were removed to avoid biasing the analysis by including duplicate samples. We used the MRB genome alignment after removal of genes with signals of recombination and hypervariable regions (1,106,409 bp with 3,047 SNPs). The timescale of *T. pallidum* molecular evolution was estimated using time-calibrated Bayesian phylogenetic inference implemented in BEAST v2.6.3 (100). The alignment was reduced to variable sites and an uncorrelated lognormal relaxed molecular clock was calibrated using the ages of the samples (see Supplementary Table 1) with a diffuse prior (uniform 0 to infinity for the mean rate and a gamma distribution (α=0.5396, β=0.3819) for the rates’ standard deviation). Historical samples, for which only age ranges (based on archaeological contexts or radiocarbon dating) rather than exact ages were available, were assigned age priors spanning uniformly across the entire range. A strict clock was rejected based on the estimated coefficient of variation for the relaxed clock model, as the 95% HPD did not include zero (101). A coalescent Bayesian Skyline tree prior with 5 groups was used as a simple model that is sufficiently flexible to fit many different kinds of dynamics. We used bModeltest v1.2.1 (102) to average across all possible reversible substitution models. According to the results, the TVM model (123421) with no gamma rate variation and no invariable sites received the most support. The MCMC chain was run for 1 billion steps with every 50,000^th^ step sampled. The first 10% of samples were discarded as burn-in. Convergence and mixing was inspected using Tracer v1.7.1 (103); the ESS of all parameters exceeded 100. The maximum clade credibility tree was generated using TreeAnnotator, a part of the BEAST v.2.6.1 software package, and visualized using FigTree v1.4.4 (http://tree.bio.ed.ac.uk/software/figtree/).

## DECLARATIONS

### Consent for publication

Not applicable

### Availability of data and materials

All data generated or analyzed during this study are included in this published article in the Supplementary Material available for this paper. All Supplementary Files are available in the following Zenodo link: https://zenodo.org/records/13375835?token=eyJhbGciOiJIUzUxMiJ9.eyJpZCI6ImE2YmY0MGMzLThmNDQtNGI4Ny04OGUwLTgzYWY4MTQ0NmVjZCIsImRhdGEiOnt9LCJyYW5kb20iOiI1N2MxYzQxNzQwMTIxNzU4MWZlZDYwODA4OTY5NWYzZCJ9.6D-_QmY2bb95pDG4ZWUA1dOXhtlEzcIRUdEmoYDjX1lLCyyOu5Vex15DKwfRKandaG19cMHC2X0a7BgATCSUBA. The newly obtained ancient genome, W86, is available under accession number ERP147184.

### Authors’ contributions

Conceptualization: VJS, FGC, KM, MPD

Data curation: MPD, GA

Formal Analysis: MPD, MM, LdP, GA

Funding acquisition: VJS, GA

Investigation: MPD, GA, KM

Methodology: FGC, MPD

Project administration: VJS, KM

Resources: HP, KD, WB, PD, MO, BK, JS, JG

Supervision: VJS, FGC, KM

Validation: FGC, KM, VJS

Visualization: MM, MPD

Writing – original draft: MPD, KM, MM, GA

Writing – review & editing: FGC, MPD, LdP, MM, VJS, NA

## Acknowledgements

We thank Anna Lipowicz from the Department of Anthropology at Wrocław University of Environmental and Life Sciences for sharing the study material.

## Authors’ information

Correspondence and requests for materials should be addressed to Verena J. Schuenemann, Kerttu Majander, or Fernando González-Candelas.

## REFERENCES

1. Noda AA, Grillová L, Lienhard R, Blanco O, Rodríguez I, Šmajs D. Bejel in Cuba: molecular identification of Treponema pallidum subsp. endemicum in patients diagnosed with venereal syphilis. Clin Microbiol Infect. 2018 Nov;24(11):1210.e1–1210.e5.

2. Grange PA, Allix-Beguec C, Chanal J, Benhaddou N, Gerhardt P, Morini JP, et al. Molecular subtyping of Treponema pallidum in Paris, France. Sex Transm Dis [Internet]. 2013; Available from: 10.1097/OLQ.0000000000000006

3. Mitjà O, Godornes C, Houinei W, Kapa A, Paru R, Abel H, et al. Re-emergence of yaws after single mass azithromycin treatment followed by targeted treatment: a longitudinal study. Lancet. 2018 Apr 21;391(10130):1599–607.

4. Elo A, Dégboé B, Barogui Y, Gomido IC, Wadagni A, d’Almeida C, et al. Resurgence of yaws in Benin: Four confirmed cases in the district of Z, Southern Benin. Journal of public health and epidemiology. 2019 Dec 31;11:201–8.

5. Giacani L, Lukehart SA. The endemic treponematoses. Clin Microbiol Rev. 2014 Jan 1;27(1):89–115.

6. Mitjà O, Šmajs D, Bassat Q. Advances in the diagnosis of endemic treponematoses: yaws, bejel, and pinta. PLoS Negl Trop Dis. 2013 Oct 24;7(10):e2283.

7. Mitjà O, Marks M, Konan DJP, Ayelo G, Gonzalez-Beiras C, Boua B, et al. Global epidemiology of yaws: a systematic review. Lancet Glob Health. 2015 Jun;3(6):e324–31.

8. Shinohara K, Furubayashi K, Kojima Y, Mori H, Komano J, Kawahata T. Clinical perspectives of Treponema pallidum subsp. Endemicum infection in adults, particularly men who have sex with men in the Kansai area, Japan: A case series. J Infect Chemother. 2022 Mar;28(3):444–50.

9. Kawahata T, Kojima Y, Furubayashi K, Shinohara K, Shimizu T, Komano J, et al. Bejel, a Nonvenereal Treponematosis, among Men Who Have Sex with Men, Japan. Emerg Infect Dis. 2019 Aug;25(8):1581–3.

10. Nishiki S, Lee K, Kanai M, Nakayama SI, Ohnishi M. Phylogenetic and genetic characterization of Treponema pallidum strains from syphilis patients in Japan by whole-genome sequence analysis from global perspectives. Sci Rep. 2021 Feb 4;11(1):3154.

11. Taouk ML, Taiaroa G, Pasricha S, Herman S, Chow EPF, Azzatto F, et al. Characterisation of Treponema pallidum lineages within the contemporary syphilis outbreak in Australia: a genomic epidemiological analysis [Internet]. Vol. 3, The Lancet Microbe. 2022. p. e417–26. Available from: 10.1016/s2666-5247(22)00035-0

12. Lieberman NAP, Lin MJ, Xie H, Shrestha L, Nguyen T, Huang ML, et al. Treponema pallidum genome sequencing from six continents reveals variability in vaccine candidate genes and dominance of Nichols clade strains in Madagascar. PLoS Negl Trop Dis. 2021 Dec;15(12):e0010063.

13. Mubemba B, Gogarten JF, Schuenemann VJ, Düx A, Lang A, Nowak K, et al. Geographically structured genomic diversity of non-human primate-infecting subsp. Microb Genom [Internet]. 2020 Nov;6(11). Available from: 10.1099/mgen.0.000463

14. Grillová L, Oppelt J, Mikalová L, Nováková M, Giacani L, Niesnerová A, et al. Directly Sequenced Genomes of Contemporary Strains of Syphilis Reveal Recombination-Driven Diversity in Genes Encoding Predicted Surface-Exposed Antigens. Front Microbiol. 2019 Jul 31;10:1691.

15. Strouhal M, Mikalová L, Haviernik J, Knauf S, Bruisten S, Noordhoek GT, et al. Complete genome sequences of two strains of Treponema pallidum subsp. pertenue from Indonesia: Modular structure of several treponemal genes. PLoS Negl Trop Dis. 2018 Oct;12(10):e0006867.

16. Mediannikov O, Fenollar F, Davoust B, Amanzougaghene N, Lepidi H, Arzouni JP, et al. Epidemic of venereal treponematosis in wild monkeys: a paradigm for syphilis origin. New Microbes New Infect. 2020 May;35:100670.

17. Timothy JWS, Beale MA, Rogers E, Zaizay Z, Halliday KE, Mulbah T, et al. Epidemiologic and Genomic Reidentification of Yaws, Liberia. Emerg Infect Dis. 2021 Apr;27(4):1123–32.

18. Liu D, Tong ML, Liu LL, Lin LR, Zhang HL, Yang TC. Characterisation of the novel clinical isolate X-4 containing a new sequence-type. Sex Transm Infect. 2021 Mar;97(2):120–5.

19. Marks M, Fookes M, Wagner J, Butcher R, Ghinai R, Sokana O, et al. Diagnostics for Yaws Eradication: Insights From Direct Next-Generation Sequencing of Cutaneous Strains of Treponema pallidum. Clin Infect Dis. 2018 Mar 5;66(6):818–24.

20. Vrbová E, Noda AA, Grillová L, Rodríguez I, Forsyth A, Oppelt J, et al. Whole genome sequences of Treponema pallidum subsp. endemicum isolated from Cuban patients: The non-clonal character of isolates suggests a persistent human infection rather than a single outbreak. PLoS Negl Trop Dis. 2022 Jun;16(6):e0009900.

21. Janečková K, Roos C, Fedrová P, Tom N, Čejková D, Lueert S, et al. The genomes of the yaws bacterium, Treponema pallidum subsp. pertenue, of nonhuman primate and human origin are not genomically distinct. PLoS Negl Trop Dis. 2023 Sep;17(9):e0011602.

22. Velasquez MR, De Lay BD, Edmondson DG, Wormser GP, Norris SJ, Cafferky K, et al. A Novel Treponema pallidum Subspecies pallidum Strain Associated With a Painful Oral Lesion Is a Member of a Potentially Emerging Nichols-Related Subgroup. Sex Transm Dis. 2024 Jul 1;51(7):486–92.

23. Yang L, Zhang X, Chen W, Seña AC, Zheng H, Jiang Y, et al. Clinical presentation of early syphilis and genomic sequences of Treponema pallidum strains in patient specimens and isolates obtained by rabbit inoculation. J Infect Dis [Internet]. 2024 Jun 17; Available from: 10.1093/infdis/jiae322

24. Schuenemann VJ, Kumar Lankapalli A, Barquera R, Nelson EA, Iraíz Hernández D, Acuña Alonzo V, et al. Historic Treponema pallidum genomes from Colonial Mexico retrieved from archaeological remains. Norris SJ, editor. PLoS Negl Trop Dis. 2018 Jun 21;12(6):e0006447.

25. Majander K, Pfrengle S, Kocher A, Neukamm J, du Plessis L, Pla-Díaz M, et al. Ancient Bacterial Genomes Reveal a High Diversity of Treponema pallidum Strains in Early Modern Europe. Curr Biol. 2020 Oct 5;30(19):3788–803.e10.

26. Giffin K, Lankapalli AK, Sabin S, Spyrou MA, Posth C, Kozakaitė J, et al. A treponemal genome from an historic plague victim supports a recent emergence of yaws and its presence in 15 century Europe. Sci Rep. 2020 Jun 11;10(1):9499.

27. Majander K, Pla-Díaz M, du Plessis L, Arora N, Filippini J, Pezo-Lanfranco L, et al. Redefining the treponemal history through pre-Columbian genomes from Brazil. Nature [Internet]. 2024 Jan 24; Available from: 10.1038/s41586-023-06965-x

28. Edmondson DG, Delay BD, Kowis LE, Norris SJ. Parameters affecting continuous in vitro culture of treponema pallidum strains. MBio. 2021 Feb 23;12(1):1–21.

29. Edmondson DG, De Lay BD, Hanson BM, Kowis LE, Norris SJ. Clonal isolates of Treponema pallidum subsp. pallidum Nichols provide evidence for the occurrence of microevolution during experimental rabbit infection and in vitro culture. PLoS One. 2023 Mar 14;18(3):e0281187.

30. Luhmann N, Doerr D, Chauve C. Comparative scaffolding and gap filling of ancient bacterial genomes applied to two ancient Yersinia pestis genomes. Microb Genom. 2017 Sep;3(9):e000123.

31. Rasmussen S, Allentoft ME, Nielsen K, Orlando L, Sikora M, Sjögren KG, et al. Early divergent strains of Yersinia pestis in Eurasia 5,000 years ago. Cell. 2015 Oct 22;163(3):571–82.

32. Pisarenko SV, Evchenko AY, Kovalev DA, Evchenko YМ, Bobrysheva OV, Shapakov NA, et al. Yersinia pestis strains isolated in natural plague foci of Caucasus and Transcaucasia in the context of the global evolution of species. Genomics. 2021 Jul;113(4):1952–61.

33. Krause-Kyora B, Nutsua M, Boehme L, Pierini F, Pedersen DD, Kornell SC, et al. Ancient DNA study reveals HLA susceptibility locus for leprosy in medieval Europeans. Nat Commun. 2018 May 1;9(1):1569.

34. Krause-Kyora B, Susat J, Key FM, Kühnert D, Bosse E, Immel A, et al. Neolithic and medieval virus genomes reveal complex evolution of hepatitis B. Elife [Internet]. 2018 May 10;7. Available from: 10.7554/eLife.36666

35. Schuenemann VJ, Singh P, Mendum TA, Krause-Kyora B, Jäger G, Bos KI, et al. Genome-wide comparison of medieval and modern Mycobacterium leprae. Science. 2013 Jul 12;341(6142):179–83.

36. Peltzer A, Jäger G, Herbig A, Seitz A, Kniep C, Krause J, et al. EAGER: efficient ancient genome reconstruction. Genome Biol. 2016 Mar 31;17:60.

37. Valiente-Mullor C, Beamud B, Ansari I, Francés-Cuesta C, García-González N, Mejía L, et al. One is not enough: On the effects of reference genome for the mapping and subsequent analyses of short-reads. PLoS Comput Biol. 2021 Jan;17(1):e1008678.

38. Staudová B, Strouhal M, Zobaníková M, Cejková D, Fulton LL, Chen L, et al. Whole genome sequence of the Treponema pallidum subsp. endemicum strain Bosnia A: the genome is related to yaws treponemes but contains few loci similar to syphilis treponemes. PLoS Negl Trop Dis. 2014 Nov 6;8(11):e3261.

39. Beale MA, Marks M, Cole MJ, Lee MK, Pitt R, Ruis C, et al. Global phylogeny of Treponema pallidum lineages reveals recent expansion and spread of contemporary syphilis. Nat Microbiol. 2021 Dec;6(12):1549–60.

40. Pla-Díaz M, Sánchez-Busó L, Giacani L, Šmajs D, Bosshard PP, Bagheri HC, et al. Evolutionary Processes in the Emergence and Recent Spread of the Syphilis Agent, Treponema pallidum. Mol Biol Evol [Internet]. 2022 Jan 7;39(1). Available from: 10.1093/molbev/msab318

41. Pankiewicz A, Witkowski J. Dewocjonalia barokowe odkryte na cmentarzysku przy kościele św. Piotra i Pawła na Ostrowie Tumskim we Wrocławiu, Wroclavia antiqua. 2012;17:1621–70.

42. Arora N, Schuenemann VJ, Jäger G, Peltzer A, Seitz A, Herbig A, et al. Origin of modern syphilis and emergence of a pandemic Treponema pallidum cluster. Nat Microbiol. 2016 Dec 5;2:16245.

43. Hodges E, Rooks M, Xuan Z, Bhattacharjee A, Benjamin Gordon D, Brizuela L, et al. Hybrid selection of discrete genomic intervals on custom-designed microarrays for massively parallel sequencing. Nat Protoc. 2009 May 28;4(6):960–74.

44. Briggs AW, Stenzel U, Johnson PLF, Green RE, Kelso J, Prüfer K, et al. Patterns of damage in genomic DNA sequences from a Neandertal. Proc Natl Acad Sci U S A. 2007 Sep 11;104(37):14616–21.

45. Sawyer S, Krause J, Guschanski K, Savolainen V, Pääbo S. Temporal Patterns of Nucleotide Misincorporations and DNA Fragmentation in Ancient DNA [Internet]. Vol. 7, PLoS ONE. 2012. p. e34131. Available from: 10.1371/journal.pone.0034131

46. Beale MA, Marks M, Sahi SK, Tantalo LC, Nori AV, French P, et al. Genomic epidemiology of syphilis reveals independent emergence of macrolide resistance across multiple circulating lineages. Nat Commun. 2019 Dec 1;10(1):1–9.

47. Strouhal M, Mikalová L, Haviernik J, Knauf S, Bruisten S, Noordhoek GT, et al. Complete genome sequences of two strains of Treponema pallidum subsp. pertenue from Indonesia: Modular structure of several treponemal genes. Caimano MJ, editor. PLoS Negl Trop Dis. 2018 Oct 10;12(10):e0006867.

48. Grillová L, Oppelt J, Mikalová L, Nováková M, Giacani L, Niesnerová A, et al. Directly Sequenced Genomes of Contemporary Strains of Syphilis Reveal Recombination-Driven Diversity in Genes Encoding Predicted Surface-Exposed Antigens. Front Microbiol. 2019 Jul 31;10:1691.

49. Pětrošová H, Zobaníková M, Čejková D, Mikalová L, Pospíšilová P, Strouhal M, et al. Whole Genome Sequence of Treponema pallidum ssp. pallidum, Strain Mexico A, Suggests Recombination between Yaws and Syphilis Strains. PLoS Negl Trop Dis [Internet]. 2012; Available from: 10.1371/journal.pntd.0001832

50. Mikalová L, Strouhal M, Oppelt J, Grange PA, Janier M, Benhaddou N, et al. Human Treponema pallidum 11q/j isolate belongs to subsp. endemicum but contains two loci with a sequence in TP0548 and TP0488 similar to subsp. pertenue and subsp. pallidum, respectively. PLoS Negl Trop Dis [Internet]. 2017; Available from: 10.1371/journal.pntd.0005434

51. Pětrošová H, Pospíšilová P, Strouhal M, Čejková D, Zobaníková M, Mikalová L, et al. Resequencing of Treponema pallidum ssp. pallidum strains Nichols and SS14: correction of sequencing errors resulted in increased separation of syphilis treponeme subclusters. PLoS One. 2013 Sep 10;8(9):e74319.

52. Mikalová L, Janečková K, Nováková M, Strouhal M, Čejková D, Harper KN, et al. Whole genome sequence of the Treponema pallidum subsp. endemicum strain Iraq B: A subpopulation of bejel treponemes contains full-length tprF and tprG genes similar to those present in T. p. subsp. pertenue strains. Clegg SR, editor. PLoS One. 2020 Apr 1;15(4):e0230926.

53. Majander K, Pfrengle S, Kocher A, Neukamm J, du Plessis L, Pla-Díaz M, et al. Ancient Bacterial Genomes Reveal a High Diversity of Treponema pallidum Strains in Early Modern Europe. Curr Biol. 2020 Oct 5;30(19):3788–803.e10.

54. Pla-Díaz M, Sánchez-Busó L, Giacani L, Šmajs D, Bosshard PP, Bagheri HC, et al. Evolutionary Processes in the Emergence and Recent Spread of the Syphilis Agent, Treponema pallidum. Mol Biol Evol [Internet]. 2022 Jan 7;39(1). Available from: 10.1093/molbev/msab318

55. Lieberman NAP, Lin MJ, Xie H, Shrestha L, Nguyen T, Huang ML, et al. Treponema pallidum genome sequencing from six continents reveals variability in vaccine candidate genes and dominance of Nichols clade strains in Madagascar. PLoS Negl Trop Dis. 2021 Dec;15(12):e0010063.

56. Forst J, Brown TA. Inability of “Whole Genome Amplification” to Improve Success Rates for the Biomolecular Detection of Tuberculosis in Archaeological Samples. PLoS One. 2016 Sep 21;11(9):e0163031.

57. Grillová L, Oppelt J, Mikalová L, Nováková M, Giacani L, Niesnerová A, et al. Directly Sequenced Genomes of Contemporary Strains of Syphilis Reveal Recombination-Driven Diversity in Genes Encoding Predicted Surface-Exposed Antigens. Front Microbiol. 2019 Jul 31;10:1691.

58. Štaudová B, Strouhal M, Zobaníková M, Čejková D, Fulton LL, Chen L, et al. Whole Genome Sequence of the Treponema pallidum subsp. endemicum Strain Bosnia A: The Genome Is Related to Yaws Treponemes but Contains Few Loci Similar to Syphilis Treponemes. Yang R, editor. PLoS Negl Trop Dis. 2014 Nov 6;8(11):e3261.

59. Addetia A, Lin MJ, Phung Q, Xie H, Huang ML, Ciccarese G, et al. Estimation of Full-Length TprK Diversity in Treponema pallidum subsp. MBio [Internet]. 2020 Oct 27;11(5). Available from: 10.1128/mBio.02726-20

60. Giacani L, Chattopadhyay S, Centurion-Lara A, Jeffrey BM, Le HT, Molini BJ, et al. Footprint of positive selection in Treponema pallidum subsp. pallidum genome sequences suggests adaptive microevolution of the syphilis pathogen. PLoS Negl Trop Dis. 2012 Jun 12;6(6):e1698.

61. Maděránková D, Mikalová L, Strouhal M, Vadják Š, Kuklová I, Pospíšilová P, et al. Identification of positively selected genes in human pathogenic treponemes: Syphilis-, yaws-, and bejel-causing strains differ in sets of genes showing adaptive evolution. PLoS Negl Trop Dis. 2019 Jun;13(6):e0007463.

62. Kumar S, Caimano MJ, Anand A, Dey A, Hawley KL, LeDoyt ME, et al. Sequence Variation of Rare Outer Membrane Protein β-Barrel Domains in Clinical Strains Provides Insights into the Evolution of subsp., the Syphilis Spirochete. MBio [Internet]. 2018 Jun 12;9(3). Available from: 10.1128/mBio.01006-18

63. Sánchez-Busó L, Comas I, Jorques G, González-Candelas F. Recombination drives genome evolution in outbreak-related Legionella pneumophila isolates. Nat Genet [Internet]. 2014;46(11). Available from: 10.1038/ng.3114

64. Pla-Díaz M, Sánchez-Busó L, Giacani L, Šmajs D, Bosshard PP, Bagheri HC, et al. Evolutionary Processes in the Emergence and Recent Spread of the Syphilis Agent, Treponema pallidum. Mol Biol Evol [Internet]. 2022 Jan 7;39(1). Available from: 10.1093/molbev/msab318

65. Beamud B, Bracho MA, González-Candelas F. Characterization of New Recombinant Forms of HIV-1 From the Comunitat Valenciana (Spain) by Phylogenetic Incongruence. Front Microbiol. 2019 May 22;10:1006.

66. Francés-Cuesta C, Ansari I, Fernández-Garayzábal JF, Gibello A, González-Candelas F. Comparative genomics and evolutionary analysis of Lactococcus garvieae isolated from human endocarditis. Microb Genom [Internet]. 2022 Feb;8(2). Available from: 10.1099/mgen.0.000771

67. Mejía L, Prado B, Cárdenas P, Trueba G, González-Candelas F. The impact of genetic recombination on pathogenic Leptospira. Infect Genet Evol. 2022 Aug;102:105313.

68. Majander K, Pfrengle S, Kocher A, Neukamm J, du Plessis L, Pla-Díaz M, et al. Ancient Bacterial Genomes Reveal a High Diversity of Treponema pallidum Strains in Early Modern Europe. Curr Biol. 2020 Oct 5;30(19):3788–803.e10.

69. Giffin K, Lankapalli AK, Sabin S, Spyrou MA, Posth C, Kozakaitė J, et al. A treponemal genome from an historic plague victim supports a recent emergence of yaws and its presence in 15 century Europe. Sci Rep. 2020 Jun 11;10(1):9499.

70. Comas I, Chakravartti J, Small PM, Galagan J, Niemann S, Kremer K, et al. Human T cell epitopes of Mycobacterium tuberculosis are evolutionarily hyperconserved. Nat Genet. 2010 Jun;42(6):498–503.

71. Harrison LB, Kapur V, Behr MA. An imputed ancestral reference genome for the Mycobacterium tuberculosis complex better captures structural genomic diversity for reference-based alignment workflows. Microb Genom [Internet]. 2024 Jan;10(1). Available from: 10.1099/mgen.0.001165

72. Dabney J, Knapp M, Glocke I, Gansauge MT, Weihmann A, Nickel B, et al. Complete mitochondrial genome sequence of a Middle Pleistocene cave bear reconstructed from ultrashort DNA fragments [Internet]. Vol. 110, Proceedings of the National Academy of Sciences. 2013. p. 15758–63. Available from: 10.1073/pnas.1314445110

73. Kircher M, Sawyer S, Meyer M. Double indexing overcomes inaccuracies in multiplex sequencing on the Illumina platform. Nucleic Acids Res. 2012 Jan;40(1):e3.

74. Meyer M, Kircher M. Illumina sequencing library preparation for highly multiplexed target capture and sequencing. Cold Spring Harb Protoc. 2010 Jun;2010(6):db.prot5448.

75. Cooper A. Ancient DNA: Do It Right or Not at All [Internet]. Vol. 289, Science. 2000. p. 1139b – 1139. Available from: 10.1126/science.289.5482.1139b

76. Briggs AW, Stenzel U, Meyer M, Krause J, Kircher M, Pääbo S. Removal of deaminated cytosines and detection of in vivo methylation in ancient DNA. Nucleic Acids Res. 2010 Apr;38(6):e87.

77. Schubert M, Lindgreen S, Orlando L. AdapterRemoval v2: rapid adapter trimming, identification, and read merging. BMC Res Notes. 2016 Feb 12;9:88.

78. Barquera R, Lamnidis TC, Lankapalli AK, Kocher A, Hernández-Zaragoza DI, Nelson EA, et al. Origin and Health Status of First-Generation Africans from Early Colonial Mexico. Curr Biol. 2020 Jun 8;30(11):2078–91.e11.

79. Yuan D, Ahamed A, Burgin J, Cummins C, Devraj R, Gueye K, et al. The European Nucleotide Archive in 2023. Nucleic Acids Res. 2024 Jan 5;52(D1):D92–7.

80. Sayers EW, Beck J, Bolton EE, Bourexis D, Brister JR, Canese K, et al. Database resources of the National Center for Biotechnology Information. Nucleic Acids Res. 2021 Jan 8;49(D1):D10–7.

81. Wingett SW, Andrews S. FastQ Screen: A tool for multi-genome mapping and quality control. F1000Res. 2018 Aug 24;7:1338.

82. Li H, Durbin R. Fast and accurate short read alignment with Burrows-Wheeler transform. Bioinformatics. 2009 Jul 15;25(14):1754–60.

83. Neukamm J, Peltzer A, Nieselt K. DamageProfiler: Fast damage pattern calculation for ancient DNA [Internet]. Available from: 10.1101/2020.10.01.322206

84. McKenna A, Hanna M, Banks E, Sivachenko A, Cibulskis K, Kernytsky A, et al. The Genome Analysis Toolkit: a MapReduce framework for analyzing next-generation DNA sequencing data. Genome Res. 2010 Sep;20(9):1297–303.

85. Alikhan NF, Petty NK, Ben Zakour NL, Beatson SA. BLAST Ring Image Generator (BRIG): simple prokaryote genome comparisons. BMC Genomics. 2011 Aug 8;12:402.

86. Beale MA, Noguera-Julian M, Godornes C, Casadellà M, González-Beiras C, Parera M, et al. Yaws re-emergence and bacterial drug resistance selection after mass administration of azithromycin: a genomic epidemiology investigation. The Lancet Microbe. 2020 Oct 1;1(6):e263–71.

87. Nguyen LT, Schmidt HA, von Haeseler A, Minh BQ. IQ-TREE: A Fast and Effective Stochastic Algorithm for Estimating Maximum-Likelihood Phylogenies. Mol Biol Evol. 2015 Jan 1;32(1):268–74.

88. Kozlov AM, Darriba D, Flouri T, Morel B, Stamatakis A. RAxML-NG: a fast, scalable and user-friendly tool for maximum likelihood phylogenetic inference. Bioinformatics. 2019 Nov 1;35(21):4453–5.

89. Lechner M, Findeiss S, Steiner L, Marz M, Stadler PF, Prohaska SJ. Proteinortho: detection of (co-)orthologs in large-scale analysis. BMC Bioinformatics. 2011 Apr 28;12(1):124.

90. Strimmer K, von Haeseler A. Likelihood-mapping: a simple method to visualize phylogenetic content of a sequence alignment. Proc Natl Acad Sci U S A. 1997 Jun 24;94(13):6815–9.

91. Shimodaira H, Hasegawa M. Multiple Comparisons of Log-Likelihoods with Applications to Phylogenetic Inference. Mol Biol Evol. 1999 Aug 1;16(8):1114–6.

92. Strimmer K, Rambaut A. Inferring confidence sets of possibly misspecified gene trees. Proc Biol Sci. 2002 Jan 22;269(1487):137–42.

93. Kumar S, Stecher G, Li M, Knyaz C, Tamura K. MEGA X: Molecular Evolutionary Genetics Analysis across Computing Platforms [Internet]. Vol. 35, Molecular Biology and Evolution. 2018. p. 1547–9. Available from: 10.1093/molbev/msy096

94. Pinto M, Borges V, Antelo M, Pinheiro M, Nunes A, Azevedo J, et al. Genome-scale analysis of the non-cultivable Treponema pallidum reveals extensive within-patient genetic variation. Nature Microbiology [Internet]. 2016; Available from: 10.1038/nmicrobiol.2016.190

95. Liu D, Tong ML, Lin Y, Liu LL, Lin LR, Yang TC. Insights into the genetic variation profile of tprK in Treponema pallidum during the development of natural human syphilis infection. PLoS Negl Trop Dis. 2019 Jul;13(7):e0007621.

96. Addetia A, Lin MJ, Phung Q, Xie H, Huang ML, Ciccarese G, et al. Estimation of Full-Length TprK Diversity in Treponema pallidum subsp. MBio [Internet]. 2020 Oct 27;11(5). Available from: 10.1128/mBio.02726-20

97. Yang Z, dos Reis M. Statistical Properties of the Branch-Site Test of Positive Selection [Internet]. Vol. 28, Molecular Biology and Evolution. 2011. p. 1217–28. Available from: 10.1093/molbev/msq303

98. Pond SLK, Muse SV. HyPhy: Hypothesis Testing Using Phylogenies. In: Statistical Methods in Molecular Evolution. New York: Springer-Verlag; 2005. p. 125–81.

99. Lu A, Guindon S. Performance of standard and stochastic branch-site models for detecting positive selection among coding sequences. Mol Biol Evol. 2014 Feb;31(2):484–95.

100. Bouckaert R, Vaughan TG, Barido-Sottani J, Duchêne S, Fourment M, Gavryushkina A, et al. BEAST 2.5: An advanced software platform for Bayesian evolutionary analysis. PLoS Comput Biol. 2019 Apr;15(4):e1006650.

101. Ho SYW, Duchêne S. Molecular-clock methods for estimating evolutionary rates and timescales [Internet]. Vol. 23, Molecular Ecology. 2014. p. 5947–65. Available from: 10.1111/mec.12953

102. Bouckaert RR, Drummond AJ. bModelTest: Bayesian phylogenetic site model averaging and model comparison. BMC Evol Biol. 2017 Feb 6;17(1):42.

103. Rambaut A, Drummond AJ, Xie D, Baele G, Suchard MA. Posterior Summarization in Bayesian Phylogenetics Using Tracer 1.7 [Internet]. Vol. 67, Systematic Biology. 2018. p. 901–4. Available from: 10.1093/sysbio/syy032

