## Supplementary information for "Insights into *Treponema pallidum* genomics from modern and ancient genomes using a novel mapping strategy"

*(1) Department of Environmental Sciences, University of Basel, Basel, Switzerland. (2) Unidad Mixta Infección y Salud Pública FISABIO/Universidad de Valencia-I2SysBio, Valencia, Spain. (3) CIBER in Epidemiology and Public Health, Valencia, Spain. (4) Institute of Evolutionary Medicine, University of Zurich, Zurich, Switzerland. (5) Centre of New Technologies, University of Warsaw, Warsaw, Poland (6) Museum and Institute of Zoology, Polish Academy of Sciences, Warsaw, Poland. (7) Department of Biosystems Science and Engineering, ETH Zürich, Basel, Switzerland. (8) Swiss Institute of Bioinformatics, Lausanne, Switzerland. (9) Department of Anatomy, Wrocław Medical University, Wrocław, Poland. (10) Faculty of Biotechnology and Food Sciences, Wrocław University of Environmental and Life Sciences, Wrocław, Poland. (11) Department of Anthropology, Wrocław University of Environmental and Life Sciences, Wrocław, Poland. (12) Zurich Institute of Forensic Medicine, University of Zurich, Zurich, Switzerland. (13) Department of Evolutionary Anthropology, University of Vienna, Vienna, Austria. (14) Human Evolution and Archaeological Sciences (HEAS), University of Vienna, Vienna, Austria.*

**Supplementary Notes**

**Supplementary Note 1. Skeletal material description for sample W86 (OT20.1)**

Dr Paweł Dąbrowski, dr Joanna Grzelak, dr hab. Maciej Oziembłowski

The grave marked OT20 is in fact a collection of bones belonging to several individuals (possibly four). Two cardboard boxes containing bones of the postcranial skeleton and 4 boxes containing skulls or their fragments were made available for research:

- **a) OT20.1** - skull without the lower jaw. Tooth sampled for genetic analysis and labeled W86. (Supplementary Note 1, Figure 1)
- **b) OT20.2** - skull without the lower jaw with a damaged left zygomatic arch.
- **c) OT20.3** - skull without the mandible, with both zygomatic arches damaged, with complete atrophy of the maxillary alveolar process.
- d) **OT20.4** - fragment of the skull: only the brain case preserved.

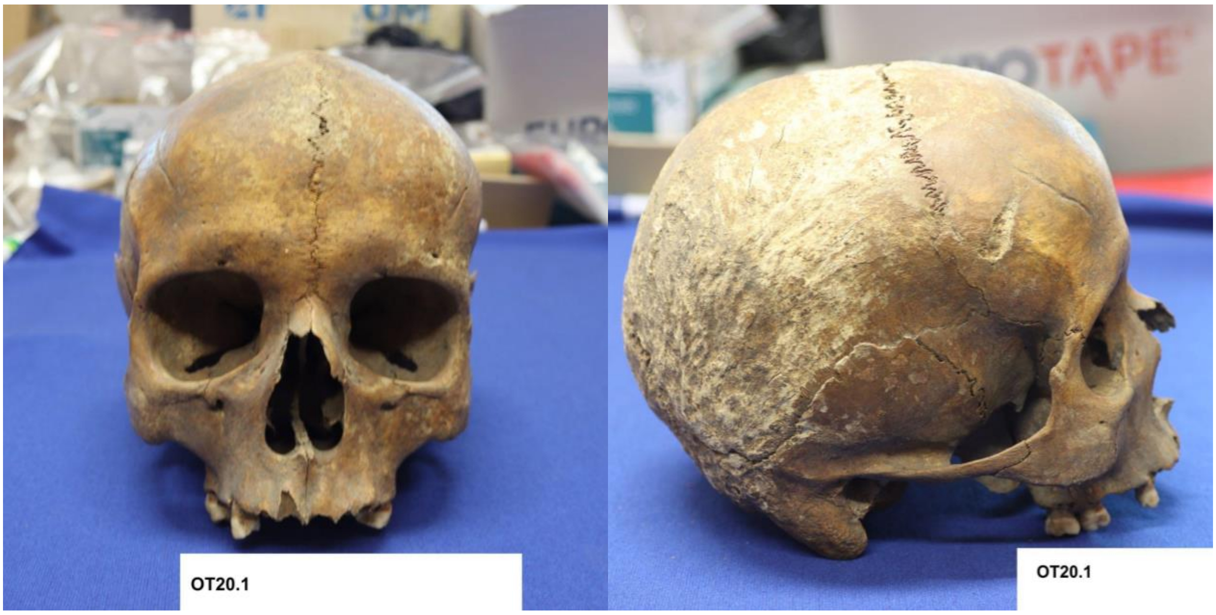

**Supplementary Note 1, Figure 1.** W86 (OT20.1) skull. a) Frontal, b) right side view. Photo: J. Grzelak

Samples for genetic testing were taken from the tooth of the subject designated as OT20.1.

The age at death was estimated on the basis of the diagnostic features of the skull: the degree of obliteration of the cranial sutures, the degree of obliteration of the spheno-occipital synchondrosis and the degree of tooth wear, as well as the condition of the metaphyses of the long bones and the condition of the ilium auricular surface of the postcranial skeleton associated to the OT20.1 skull. The age at the time of death of an OT20.1 individual is defined as juvenis / adultus (end of adolescence / beginning of adulthood). The beginning of adulthood (adultus) is considered to be the complete closure of the synchondrosis fissure. In the case under study, it is incomplete (Supplementary Note 1, Figure 1a).

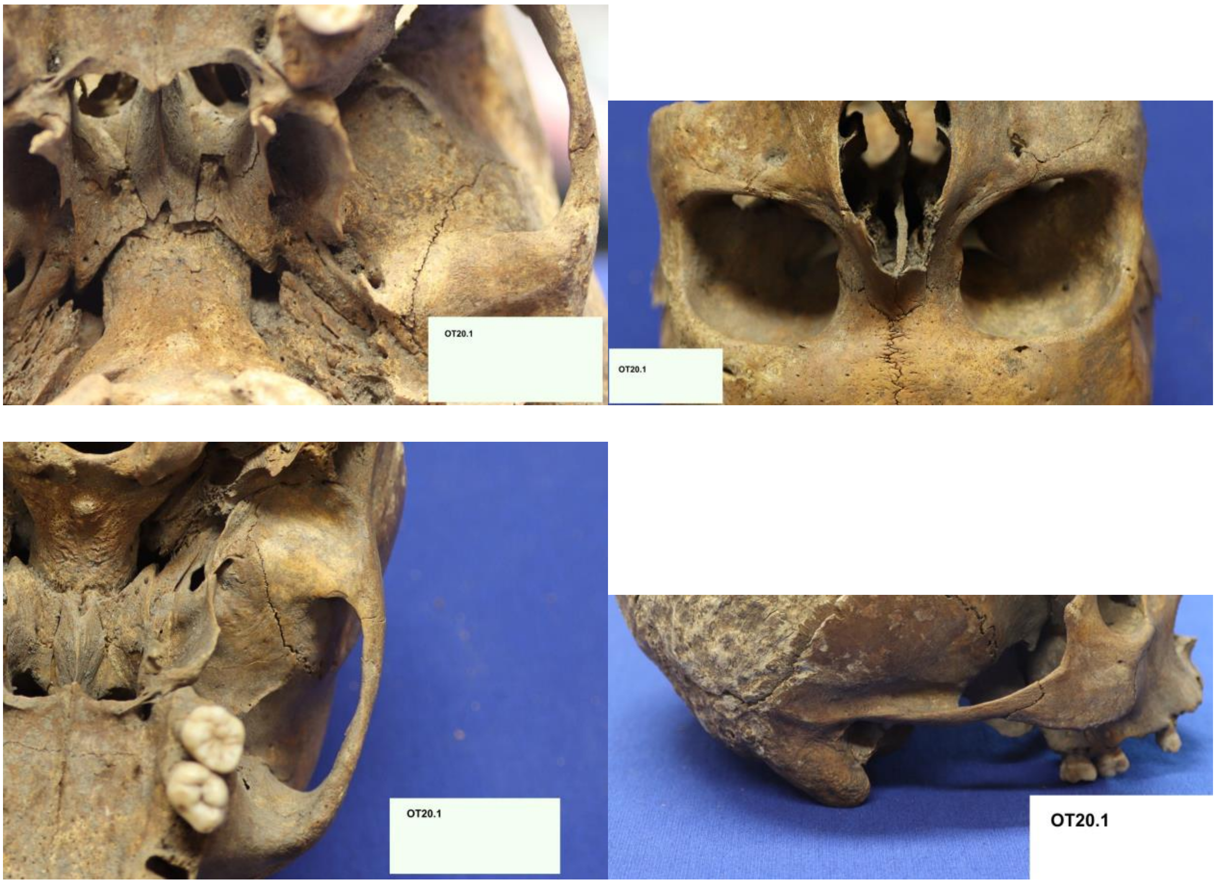

**Supplementary Note 1, Figure 2**. Morphological features of skull W86 (OT20.1). a) Incomplete closure of the synchondrosis fissure, b) thickened orbit, c) massive zygomatic arch, d) massive mastoid process. Photo: J. Grzelak

The gender of the OT20.1 subject was assessed as male. The assessment was based on the following features:

- Orbital morphology: the lateral part of the upper edge of the orbit is thickened, which results from the thickening of the superciliary arch - a characteristic of the male sex (Supplementary Note 1, Figure 2b),
- Massiveness of the zygomatic arch - the lower edge is thickened, which is related to the extensive attachment of the masseter muscle in the male sex (Supplementary Note 1, Figure 2c),
- The mastoid process is very massive, which indicates a very well-developed sternocleidomastoid muscle (Supplementary Note 1, Figure 2d).

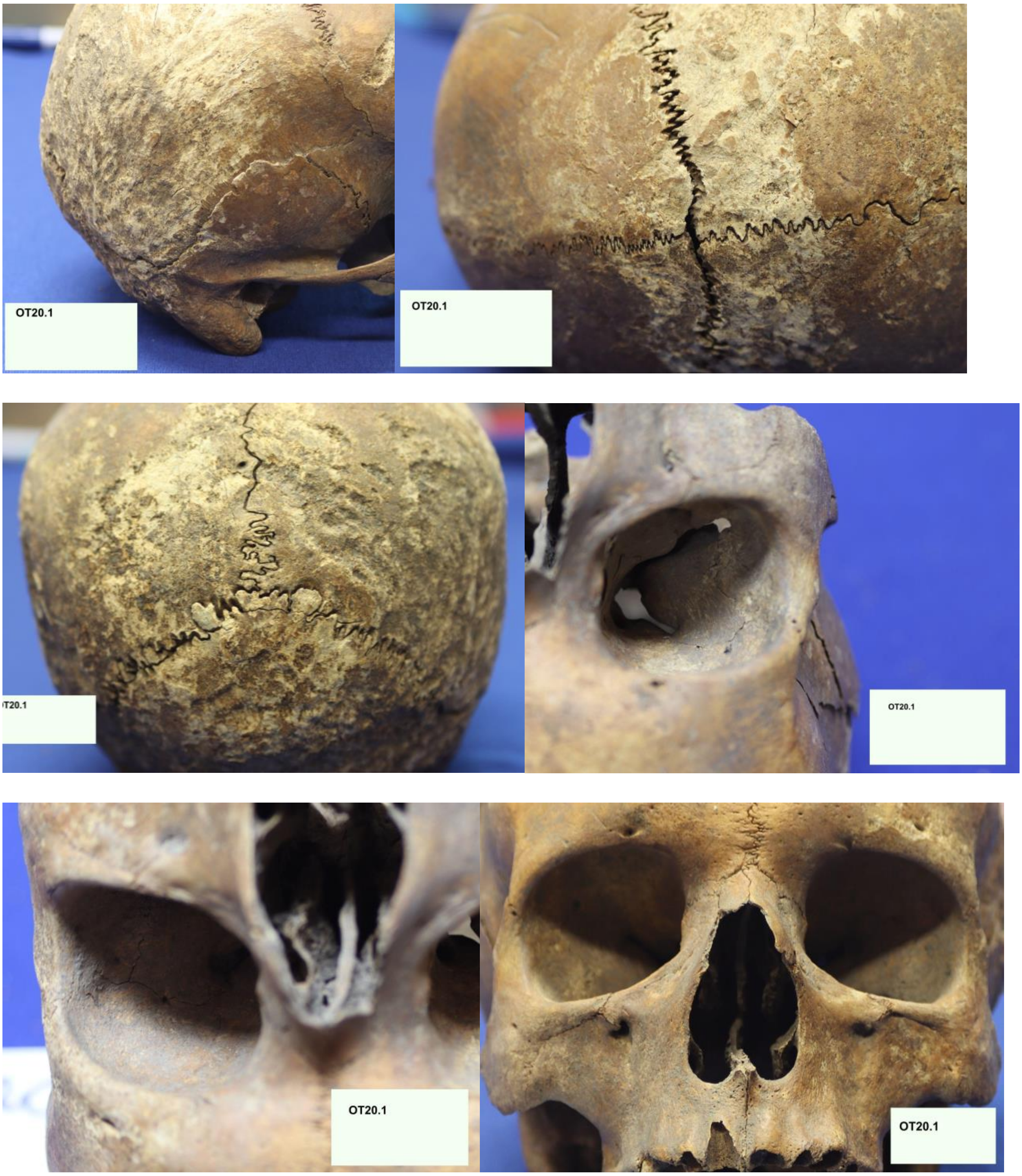

**Supplemental Note 1, Figure 3**. Paleopathological examination. a) scales of the occipital bone and the scales of the parietal bone, b) degree of cranial sutures fusion, c) coronal and lambdoid sutures, d) right orbit, e) left orbit, f) facial part of the cranium

Unfortunately, not all diagnostic features could be assessed: the outer surface of the occipital bone and the posterior part of the parietal bone turned out to be so mechanically damaged that it was impossible to describe the diagnostic features. In turn, the so-called pseudopathological changes, resulting from taphonomic processes, were observed; hence it is impossible to assess the sculpture of the scales of the occipital bone and the scales of the parietal bone (Supplemental Note 1, Figure 3a). Doubts as to gender assessment are also raised by the presence of parietal tumors and the proportions of the examined skull. They appear in the form characteristic for an individual of the female sex (Fig. 1b). It should be emphasized that the uncertainty in determining the sex may be related to the young age of the examined person and the developmental changes observed in the skull. The skull is characterized by very wide, open seams and a preserved metopic suture (Supplemental Note 1, Figure 3a and 3b). With regard to pathological changes and devupplemental Note 1, Figure 3d and 3e).

Similarly, the absence of hypoplastic defects in enamel may indicate that this individual was well nourished in early childhood. No carious lesions were found in the preserved tooth crowns of permanent teeth. No visible bone syphilis lesions at the anterior surface of the maxillary body and the edge of the nasal notch (Supplemental Note 1, Figure 3f). The superficial changes in the neurocranium do not present an image characteristic for the inflelopmental disorders, it was found that in the coronal and lambdoid sutures there are small insertion bones, which may indicate disturbances in the growth and ossification process (Supplemental Note 1, Figure 3c). No changes in the form of *cribra orbitalia* – orbital roof lesions related to deficiency anemia, were observed (Sammatory process in the diagnosis of syphilis; in our opinion, it is the result of taphonomic processes occurring between the skeleton and the filling of the burial cavity.

The sample was a part of a human genetic analysis conducted at the Museum and Institute of Zoology, Polish Academy of Sciences as a part of the University of Wrocław research project. The DNA libraries from this project were screened for presence of a range of pathogens using PCR amplification of chosen genetic markers (i.a. 106 bp *T. pallidum* arp gene fragment (1). As sample W86 (and two other samples from the project) tested positive by PCR for arp, it was sent to the Paleogenetics Laboratory at the University of Zurich for further *Treponema*-targeted analysis.

**Supplementary Note 2. Chemical analysis of sample W86 (OT20.1)**

In OT20.1, the content of calcium, phosphorus and alkaline earth elements (Sr and Ba) was determined in the bone tissue collected from the rib. Then the proportions of these elements were calculated.

The Ca / P ratio = 1.65 indicates a relatively good state of conservation of the material.

The proportions of strontium and barium to calcium are significantly higher than the average in the studied group, which may indicate a diversified diet, with the use of plant-based foods and / or dairy products.

**Supplementary Note 2, Table 1.** Proportions of Sr / Ca and Ba / Ca against the background of the studied group.

| Proportion | Average | OT20.1 | Statistical significance |
|---|---|---|---|
| Sr/Ca | -7.71 | -7.35 | P=0.000 |
| Ba/Ca | -9.05 | -7.64 | P=0.000 |

**Supplementary Note 3. Repetitive samples comparison**

For this study, we included in the new dataset generated (see Supplementary Table 1), four samples—IND1, K363, Nichols, and NIC2—originally derived from the same two clinical samples. The inclusion of these sequences aimed to discern potential variations resulting from the distinct methods employed in their acquisition. These four samples have undergone sequencing in different studies, employing diverse technologies and processing methods.

IND1, obtained through the enrichment of treponemal DNA based on the hybridization capture technique, directly originated from the original clinical sample and was featured in Arora et al. (2). In contrast, K363 was acquired using the Whole Genome Amplification (WGA) technique, as detailed by Strouhal et al. (3). K363 was first passaged in Syrian Golden Hamsters and then underwent several passages in New Zealand White Rabbits.

Nichols and NIC2 both originated from the same original sample, which underwent rabbit culture to enhance DNA content. Nichols, featured in Petrosova et al. (2), was sequenced using the Whole Genome Amplification (WGA) technique. In contrast, NIC2, featured in Arora et al. (2), was sequenced using an enrichment of treponemal DNA based on hybridization capture.

When we compared the sequence of the four samples, we obtained 86 SNPs between IND1 and K363, and 18 SNPs between Nichols and NIC2.

Regarding the placement of these four strains in the reference phylogeny, as depicted in the whole genome alignment with and without genes showing signal of recombination (Figure 2), it is noteworthy that Nichols and NIC2 maintain the same position in both trees. However, IND1 and K363 exhibit more divergent positions in the whole genome reference tree compared to the reference tree when recombination was removed. This discrepancy arises from the identification of 84 out of the 86 SNPs found between IND1 and K363, primarily located in the tRNAs *tp0012* and *tp0015*. These tRNAs were flagged as potential recombinant genes, leading to their removal from the phylogenetic tree without considering recombination (see Figure 2B). Although both genes were labeled as recombinant, we cannot completely dismiss the possibility that the SNPs in these genes may be artifacts or the result of contamination.

**Supplementary Note 4. Intermediate results of the PIM procedure: Likelihood mapping and Topology tests**

After selecting 317 genes with more than 3 SNPs present in at least one reference genome (Supplementary Table 4), a likelihood mapping (4) (LM) test was performed using IQTREE to ascertain which genes had some phylogenetic signal. Briefly, the test constructs unrooted phylogenetic trees for each quartet (subsets of four sequences) in the data. Quartet likelihoods are then mapped to a triangle, where the location represents the "treelikeness" of the quartet. Quartets in corners are fully resolved, on the sides partially resolved and in the central zone unresolved. From the 317 genes, there were 53 genes with too much missing data to perform the LM test. In order to include them in the PIM pipeline, for each individual gene, problematic sequences with more than 50% of positions with missing data were removed. Then, the Likelihood-mapping test was performed and 160 genes were retained (Supplementary Table 5) and the remaining genes were discarded, because for all quartets considered, the distribution of the corresponding likelihoods fell in the central zone of the triangle, a key graphical representation in the test used to visualize phylogenetic signal. When likelihoods cluster in this central zone, it indicates a weak phylogenetic signal, suggesting that the gene does not provide clear evolutionary information. Next, we tested the phylogenetic congruence between trees, comparing the tree obtained from each retained gene and the provisional reference tree obtained from the complete genome alignment using the SH and ELW topology tests. From the 91 genes that showed phylogenetic incongruence, 18 were further verified to contain at least three consecutive SNPs, thus supporting a recombination event (Supplementary table 6).

Subsequently, we detailed the recombination results by PIM obtained of each one of the SRB genome datasets:

Out of the 1125 genes extracted from the CDC2-SRB genome dataset according to the CDC2 genomic reference coordinates, 302 with more than 3 SNPs underwent recombination analysis through the PIM pipeline. However, extensive missingness of data limited the topology test to 230 genes, identifying 47 with reciprocal incongruence, but only 10 as recombinant genes. Those 10 genes were among those obtained with the MRB genome alignment, while the remaining 10 recombinant genes detected using the MRB genome alignment could not be tested in the CDC2-SRB genome dataset due to a significant proportion of missing data.

Similarly, from the 1122 genes extracted from the BosniaA-SRB genome dataset according to the BosniaA genomic reference coordinates, 301 genes with more than 3 SNPs were analyzed for recombination. However, due to substantial missingness of data, the topology test could only be conducted for 229 genes, revealing 46 with reciprocal incongruence, of which 11 were recombinant. Those 11 genes were among those obtained with the MRB genome alignment, while the remaining 9 recombinant genes detected using the MRB genome alignment could not be tested in the BosniaA-SRB genome dataset due to a significant proportion of missing data.

For the 1035 genes extracted from the Nichols-SRB genome dataset according to the Nichols genomic reference coordinates, 306 genes with more than 3 SNPs were analyzed for recombination. Despite substantial missingness of data, the topology test considered 238 genes, indicating 48 with reciprocal incongruence and 12 of those as recombinant. Those 12 genes were among those obtained with the MRB genome alignment, while the remaining 8 genes detected employing the MRB genome alignment could not be tested in the Nichols-SRB genome dataset due to a significant proportion of missing data.

Finally, among the 1032 genes extracted from the SS14-SRB genome dataset according to the SS14 genomic reference coordinates, 289 genes with more than 3 SNPs underwent recombination analysis. However, due to significant missingness of data, the topology test was limited to 226 genes, revealing 43 with reciprocal incongruence, of which 11 were recombinant. Those 11 genes were among those obtained with the MRB genome alignment, while the remaining 9 recombinant genes detected in the MRB genome alignment analysis could not be tested in the SS14-SRB genome dataset due to a significant proportion of missing data.

#### Supplementary Tables

**Supplementary Table 1.** Genome reconstruction details for all published samples and genomes used in the analyses. Type file: excel file

**Supplementary Table 2.** Orthology analysis results obtained by Proteinortho showing the detected orthologous genes in the four reference genomes employed. Type file: excel file

**Supplementary Table 3.** The new genomic coordinates of each gene present in at least one of the four reference genomes, calculated according to their corresponding location in the final merged alignment. The new genes were named according to the Proteinortho results obtained, calling them using the acronyms of TPASS, TPANIC, TENBA or TPECDC2 depending on the orthology funded and their presence or absence is each reference genome. Type file: excel file

**Supplementary Table 4.** The number of SNPs detected per gene present in at least one of the four different reference genomes employed in the mapping analysis. Type file: excel file

**Supplementary Table 5.** The results of the likelihood-mapping test performed. 160 genes showed some phylogenetic signal and were retained for the subsequent analyses. (Zones 1-3 represent cases in which one topology has a significantly higher likelihood than the two alternative topologies; zones 4-6 represent cases in which one topology has a significantly lower likelihood than the other two, and zone 7 represents cases in which all the topologies have similar likelihoods, hence the corresponding gene does not carry enough phylogenetic signal to differentiate between the 3 evolutionary hypotheses tested in each case.). Type file: excel file

**Supplementary Table 6.** Topology test results for the genes retained in previous PIM steps. The table presents the results of the two tests (p-values for SH, and a posterior weight for ELW). All these genes were examined in detail *a posteriori* because the two tests rejected the reference tree topology with the gene alignment ($p < 0.20$, weight value close to 0) and the complete genome alignment rejected the gene topology (reciprocal incongruence, $p < 0.05$ and weight value close to 0). Type file: excel file

**Supplementary Table 7.** Number of SNPs for each gene extracted from the alignment of each genome-mapping dataset, based on the coordinates of those genes in the respective genomic referenceType file: excel file

**Supplementary Table 8.** Topology test results for genes retained in previous PIM steps, using four different reference genomes in the mapping analysis. The tables display p-values for the SH test and posterior weights for ELW. Genes were examined further if both tests rejected the reference tree topology with the gene alignment ($p < 0.20$, weight value close to 0), and the complete genome alignment rejected the gene topology (reciprocal incongruence, $p < 0.05$, and weight value close to 0). Type file: excel file

**Supplementary Table 9.** Results of the aBSREL test, a "branch-site" model implemented in HyPhy to study the effects of positive selection along all different lineages on the phylogeny of the 18 putative recombinant genes (Table 1) on the phylogeny of genes with 3 or more SNPs present in at least one reference genome. The total of branches tested and the number of branches under positive selection are indicated in the table plus the strains inside in each node or branch and the p-value obtained. The color legend is detailed below. Type file: excel file

**Supplementary Table 10.** Functional significance of the genes detected as recombinant and/or under positive selection according to Uniprot or in the literature. Type file: excel file

#### Supplementary Figures

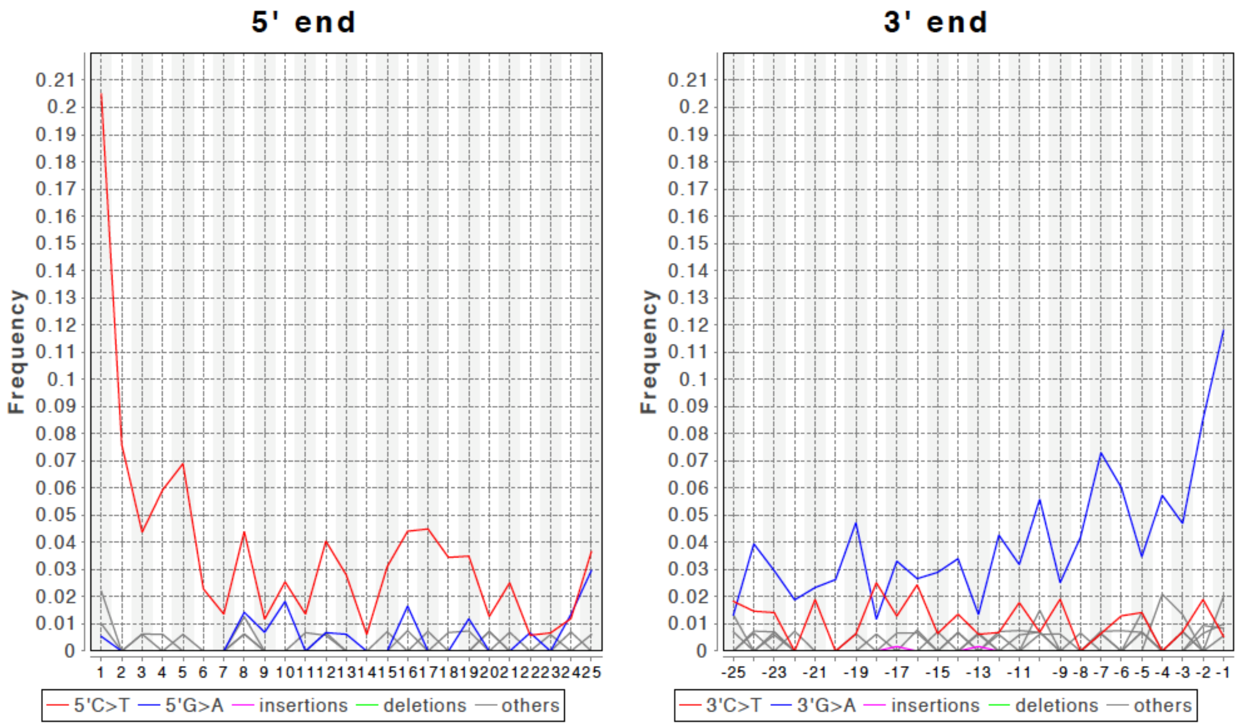

**Supplementary Figure 1**. Damage profile obtained by MapDamage program showing the misincorporation patterns and the damage at the end of sequencing reads of the new historical W86 genome obtained before the capture process. A pattern of cytosine-to-thymine base misincorporation accumulated at the end of the reads is indicative of authentic ancient DNA in the sample.

**Supplementary Figure 2.** A) Maximum likelihood tree (ML) obtained by IQ-TREE using GTR+G+I as the evolutionary model with 1000 bootstrap replicates using the final whole-genome alignment generated. B) Maximum likelihood tree (ML) obtained by IQ-TREE using GTR+G+I as the evolutionary model with 1000 bootstrap replicates using the CDC2-mapping genome alignment generated. Type file: pdf file

**Supplementary Figure 3.** A) Maximum likelihood tree (ML) obtained by IQ-TREE using GTR+G+I as the evolutionary model with 1000 bootstrap replicates using the final whole-genome alignment generated. B) Maximum likelihood tree (ML) obtained by IQ-TREE using GTR+G+I as the evolutionary model with 1000 bootstrap replicates using the BosniaA-mapping genome alignment generated. Type file: pdf file

**Supplementary Figure 4.** A) Maximum likelihood tree (ML) obtained by IQ-TREE using GTR+G+I as the evolutionary model with 1000 bootstrap replicates using the final whole-genome alignment generated. B) Maximum likelihood tree (ML) obtained by IQ-TREE using GTR+G+I as the evolutionary model with 1000 bootstrap replicates using the Nichols-mapping genome alignment generated. Type file: pdf file

**Supplementary Figure 5.** A) Maximum likelihood tree (ML) obtained by IQ-TREE using GTR+G+I as the evolutionary model with 1000 bootstrap replicates using the final whole-genome alignment generated. B) Maximum likelihood tree (ML) obtained by IQ-TREE using GTR+G+I as the evolutionary model with 1000 bootstrap replicates using the SS14-mapping genome alignment generated. Type file: pdf file

**Supplementary Figure 6.** Maximum likelihood tree obtained with all genes included in the whole-genome alignment without collapse SS14-Ω. The different clades corresponding to yaws (TPE) and bejel (TEN) subspecies, and the Nichols and the different SS14 lineages of the syphilis clade (TPA) are indicated in the figure with colors, according to the corresponding color legend. Bootstrap support values higher than 70% are indicated by red circles, which are larger when they are better supported nodes.

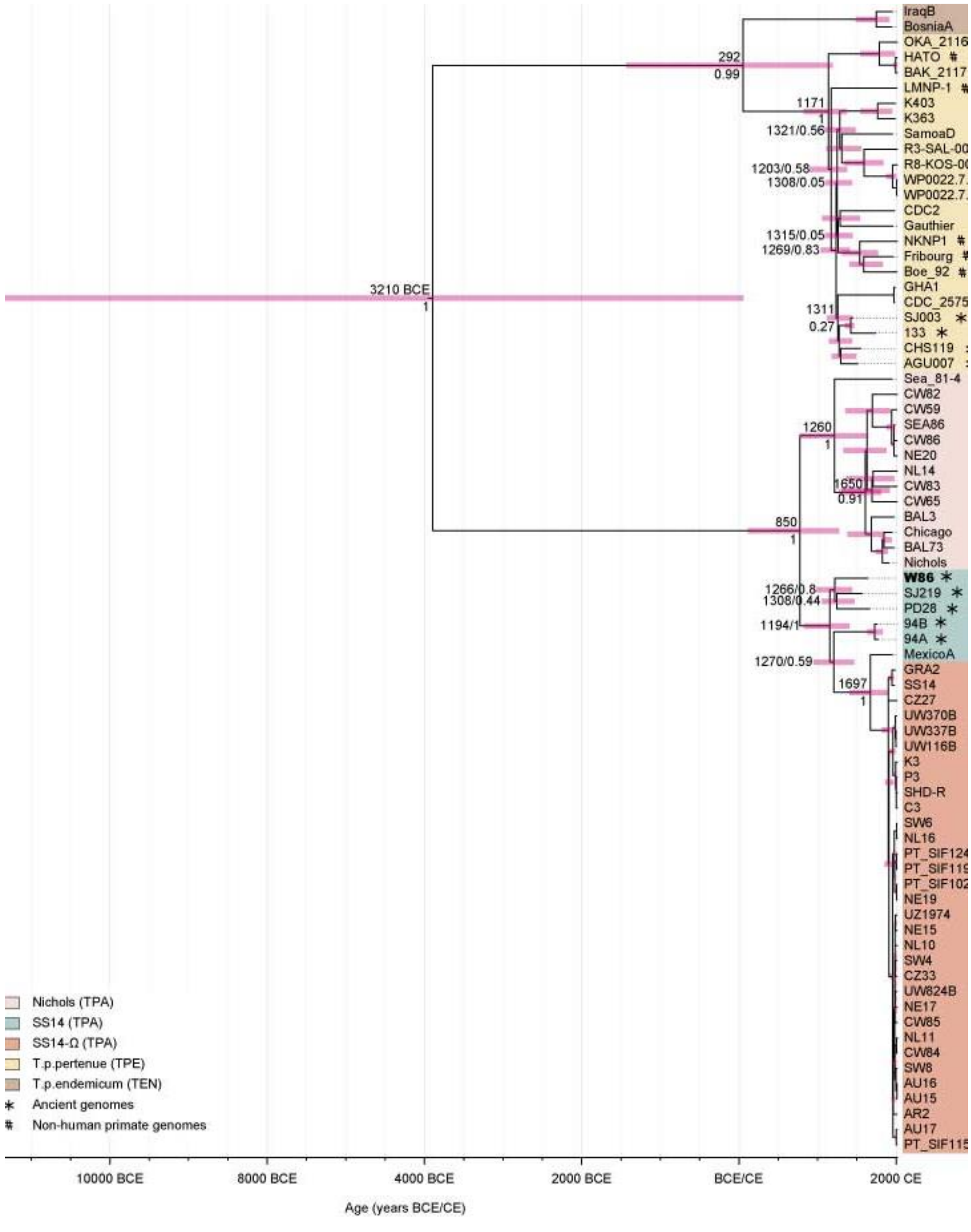
IraqB
BosniaA
OKA_2116
HATO #
BAK_2117
LMNP-1 #
K403
K363
SamoaD
R3-SAL-00
R8-KOS-00
WP0022.7
WP0022.7
CDC2
Gauthier
NKNP1 #
Fribourg #
Boe_92 #
GHA1
CDC_2575
SJ003 *
133 *
CHS119
AGU007
Sea_81-4
CW82
CW59
SEA86
CW86
NE20
NL14
CW83
CW65
BAL3
Chicago
BAL73
Nichols
W86 *
SJ219 *
PD28 *
94B *
94A *
MexicoA
GRA2
SS14
CZ27
UW370B
UW337B
UW116B
K3
P3
SHD-R
C3
SW6
NL16
PT_SIF124
PT_SIF119
PT_SIF102
NE19
UZ1974
NE15
NL10
SW4
CZ33
UW824B
NE17
CW85
NL11
CW84
SW8
AU16
AU15
AR2
AU17
PT_SIF115
292
0.99
1171
1
1321/0.56
1203/0.58
1308/0.05
1315/0.05
1269/0.83
3210 BCE
1
1311
0.27
1260
1
1650
0.91
850
1
1266/0.8
1308/0.44
1194/1
1270/0.59
1697
1
Nichols (TPA)
SS14 (TPA)
SS14-Ω (TPA)
T.p.pertenue (TPE)
T.p.endemicum (TEN)
* Ancient genomes
### Non-human primate genomes
10000 BCE
8000 BCE
4000 BCE
2000 BCE
BCE/CE
2000 CE
Age (years BCE/CE)

**Supplementary Figure 7**. Maximum clade credibility (MCC) tree of the dataset consisting of 66 modern (IND1 and NIC2 samples removed) and 9 ancient genomes (newly sequenced sample, W86, has been bolded) with tp0897, tp0316 and tp0317 and recombinant genes excluded, estimated in BEAST2 v2.6.3 under an uncorrelated lognormal relaxed clock model and a Bayesian skyline plot demographic model. Median age and Posterior Bayesian support estimates are provided for selected nodes. Pink bars show node age 95% HPD. Type file: pdf file

#### Supplementary Files

All Supplementary Files bellow are available in the next Zenodo link: https://zenodo.org/records/13375835?token=eyJhbGciOiJIUzUxMiJ9.eyJpZCI6ImE2YmY0MGMzLThmNDQtNGI4Ny04OGUwLTgzYWY4MTQ0NmVjZCIsImRhdGEiOnt9LCJyYW5kb20iOiI1N2MxYzQxNzQwMTIxNzU4MWZlZDYwODA4OTY5NWYzZCJ9.6D-_QmY2bb95pDG4ZWUA1dOXhtlEzcIRUdEmoYDjX1lLCyyOu5Vex15DKwfRKandaG19cMHC2X0a7BgATCSUBA

**Supplementary File 1.** Mapping-genome alignment of the 77 *T. pallidum* genomes using CDC2 genome as reference. Type file: fasta file

**Supplementary File 2.** Mapping-genome alignment of the 77 *T. pallidum* genomes using BosniaA genome as reference. Type file: fasta file

**Supplementary File 3.** Mapping-genome alignment of the 77 *T. pallidum* genomes using SS14 genome as reference. Type file: fasta file

**Supplementary File 4. M**apping-genome alignment of the 77 *T. pallidum* genomes using Nichols genome as reference. Type file: fasta file

**Supplementary File 5.** Multiple sequence alignment of 77 *T. pallidum* genomes using the MRB strategy. Type file: fasta file

**Supplementary File 6.** Multiple sequence alignment of 77 *T. pallidum* genomes using the MRB strategy after removing the 20 recombinant genes along with the *tp0316*, *tp0317* and *tp0897* genes from the multiple genome alignment. Type file: fasta file

**Supplementary File 7.** In-house script A to ensure that the final Multiple sequence alignment obtained was correct. Type file: fasta file

**Supplementary File 8.** In-house script B to ensure that the final Multiple sequence alignment obtained was correct. Type file: fasta file

**Supplementary File 9.** In-house script to compute the number of SNPs per gene. Type file: fasta file

#### References

1. Montiel R, Solórzano E, Díaz N, Álvarez-Sandoval BA, González-Ruiz M, Cañadas MP, et al. Neonate human remains: a window of opportunity to the molecular study of ancient syphilis. PLoS One. 2012 May 2;7(5):e36371.

2. Arora N, Schuenemann VJ, Jäger G, Peltzer A, Seitz A, Herbig A, et al. Origin of modern syphilis and emergence of a pandemic Treponema pallidum cluster. Nat Microbiol. 2016 Dec 5;2:16245.

3. Strouhal M, Mikalová L, Haviernik J, Knauf S, Bruisten S, Noordhoek GT, et al. Complete genome sequences of two strains of Treponema pallidum subsp. pertenue from Indonesia: Modular structure of several treponemal genes. Caimano MJ, editor. PLoS Negl Trop Dis. 2018 Oct 10;12(10):e0006867.

4. Strimmer K, von Haeseler A. Likelihood-mapping: a simple method to visualize phylogenetic content of a sequence alignment. Proc Natl Acad Sci U S A. 1997 Jun 24;94(13):6815–9.
